# Fine-scale population structure reveals high genetic heterogeneity of the Kuwaiti population in the Arabian Peninsula

**DOI:** 10.1101/2020.11.23.393892

**Authors:** Muthukrishnan Eaaswarkhanth, Ajai K Pathak, Linda Ongaro, Francesco Montinaro, Prashantha Hebbar, Osama Alsmadi, Mait Metspalu, Fahd Al-Mulla, Thangavel Alphonse Thanaraj

## Abstract

Recent studies have showed the diverse genetic architecture of the highly consanguineous populations inhabiting the Arabian Peninsula. Consanguinity coupled with heterogeneity is complex and makes it difficult to understand the bases of population-specific genetic diseases in the region. Therefore, comprehensive genetic characterization of the populations at the finest scale is warranted. Here, we revisit the genetic structure of the Kuwait population by analyzing genome-wide single nucleotide polymorphisms data from 583 Kuwaiti individuals sorted into three subgroups. We envisage a diverse demographic genetic history among the three subgroups based on drift and allelic sharing with modern and ancient individuals. Furthermore, our comprehensive haplotype-based analyses disclose a high genetic heterogeneity among the Kuwaiti populations. We infer the major sources of ancestry within the newly defined groups; one with an obvious predominance of sub-Saharan/Western Africa mostly comprising Kuwait-B individuals, and other with West Eurasia including Kuwait-P and Kuwait-S individuals. Overall, our results recapitulate the historical population movements and reaffirm the genetic imprints of the legacy of continental trading in the region. Such deciphering of fine-scale population structure and their regional genetic heterogeneity would provide clues to the uncharted areas of disease-gene discovery and related associations in populations inhabiting the Arabian Peninsula.

## Introduction

The Arabian Peninsula is a melting pot of human diversity and culture. Recently emerging archaeological findings and ancient artifacts indicate that humans occupied the Arabian Peninsula much earlier than previously thought, thereby providing a new paradigm of the migration of modern humans from Africa [1,2]. The geographical location of the Arabian Peninsula, as a land bridge connecting Africa and Eurasia, was a gateway for human migration from the African continent to the rest of the world [3]. Modern humans not only migrated through the region but also populated the Arabian Peninsula and established distinct settlements [4–6]. The early nomadic lifestyle in the arid environment, in addition to the frequent influx of other populations, subsequent admixture, and consanguineous practices drove the emergence of indigenous ethnic populations throughout the Arabian Peninsula [5]. The multiple waves of migration of populations from Europe, Asia, and Africa drove the genetic diversity of ethnic Arab populations. For example, human populations that migrated to Qatar have been genetically categorized into three groups based on putative ancestors from Arabia/Bedouin, Persia/South Asia, and Africa [7,8]. As evident from a few genetic studies, genetically heterogeneous groups currently populate the Arabian Peninsula [9,10]. Our recent genome-wide selection analysis also highlights this inter-regional genetic heterogeneity [11]. The prevalence of genetic heterogeneity in the highly consanguineous Arab populations increases the complexity of understanding the genetic bases and etiology of various Arab-specific diseases. Notably, more than 1100 different genetic diseases have been reported in Arab populations, of which 60% are autosomal recessive and 44% are restricted to a specific population or geographical region [12]. Hence, there is an urgent need for the genetic characterization of these regional populations at the finest level. Therefore, it is imperative to comprehend the fine-scale genetic structures and regional heterogeneities of populations inhabiting the Arabian Peninsula. However, the details surrounding this context remain understudied. In consideration of this background, we conducted high-resolution population genetic analyses to better understand the fine-scale patterns of the genetic diversity and the extent of genetic heterogeneity of the Arab people of Kuwait.

Kuwait is located in the northeastern corner of the Arabian Peninsula and is bordered by Saudi Arabia to the south and Iraq to the north. The State of Kuwait was established during the early 18^th^ century by pastoral-nomadic tribes that migrated from Saudi Arabia [13] and underwent a dramatic transformation after the discovery of oil in the late 19^th^ century [14]. Apart from cattle ranching, the early settlers were involved in various occupations such as fishing and merchant seafaring for survival [15]. The large-scale maritime trade activities with India and Africa set Kuwait as a nexus connecting India, Arabia, and Persia to Europe [14]. These occupations stratified groups socially into aristocrats, merchants, and nomads (Bedouins) [16]. The oil boom endowed Kuwait with an affluent economy, high employment opportunities, migration of skilled workers from Asia and Africa, high-tech infrastructures, and westernized lifestyles. The resettlement of peoples from neighboring regions, mainly Saudi Arabia and Persia, and the admixture of various populations, facilitated gene flow [17], thereby increasing the genetic diversity. As Arab populations are well-known for their overwhelming consanguinity, Kuwait is no exception and will eventually face the consequences in the form of severe recessive diseases [18]. Studies based on uniparental markers have delineated the maternal [19-21] and paternal [22,23] diversity in Kuwait. An earlier study based on genome-wide single nucleotide polymorphisms (SNPs) conducted by our group delineated the genetic substructure of the Kuwait population based on their surnames and genetic ancestry [24]. Accordingly, putative ancestors of the current Kuwaiti population can be traced to Saudi Arabian (Kuwait-S), Persian (Kuwait-P), and Bedouin (Kuwait-B) populations, as further corroborated by whole exome- and genome-based studies [25-28]. However, the extent of genetic heterogeneity and fine-scale population structure have not yet been elucidated. Therefore, we conducted the present investigation to expand the findings of our previous studies by considerably increasing the sample size and comparing them with the growing body of genomes from modern and even ancient specimens across West Eurasia, in addition to applying multiple haplotype-based analyses on a genome-wide scale.

## Materials and methods

### Samples

We included samples of 620 Kuwaiti individuals from the State of Kuwait who were part of a larger cohort collected for studies of metabolic disorders [24,29,30]. All participants were healthy and enrolled after obtaining written informed consent. The study protocol was approved by the Ethical Review Committee of the Dasman Diabetes Institute (Kuwait City, Kuwait). Participant recruitment, sample collection, and related procedures were conducted in accordance with the tenets of the Declaration of Helsinki, as detailed elsewhere [24,29,30]. Each of the 620 individuals were assigned to subgroups (Kuwait-P, Kuwait-S and Kuwait-B) as described elsewhere [24].

### Genotyping and quality control

All the 620 individuals were genotyped using HumanOmniExpress arrays for 730,525 SNPs (Illumina, Inc., San Diego, CA, USA) in accordance with the manufacturers’ protocol. Quality control checks and data filtering were conducted using the PLINK (version 1.9) whole genome data analysis toolset [31]. The dataset was filtered to contain only autosomal SNPs with a genotyping success rate of >90%, a minor allele frequency of >5%, and a probability (*p*) value of >0.001 as determined using the Hardy-Weinberg exact test. Only samples with a genotyping success rate of >90% were included for analysis. In addition, we estimated pairwise identity-by-descent (IBD) with a threshold of IBD proportion > 0.125 and randomly removed one sample per pair of related individuals. After QC and relatedness filtering, 583 individuals and 587,819 SNPs met the inclusion criteria for analysis (Supplementary Fig. 1).

**Fig. 1.**
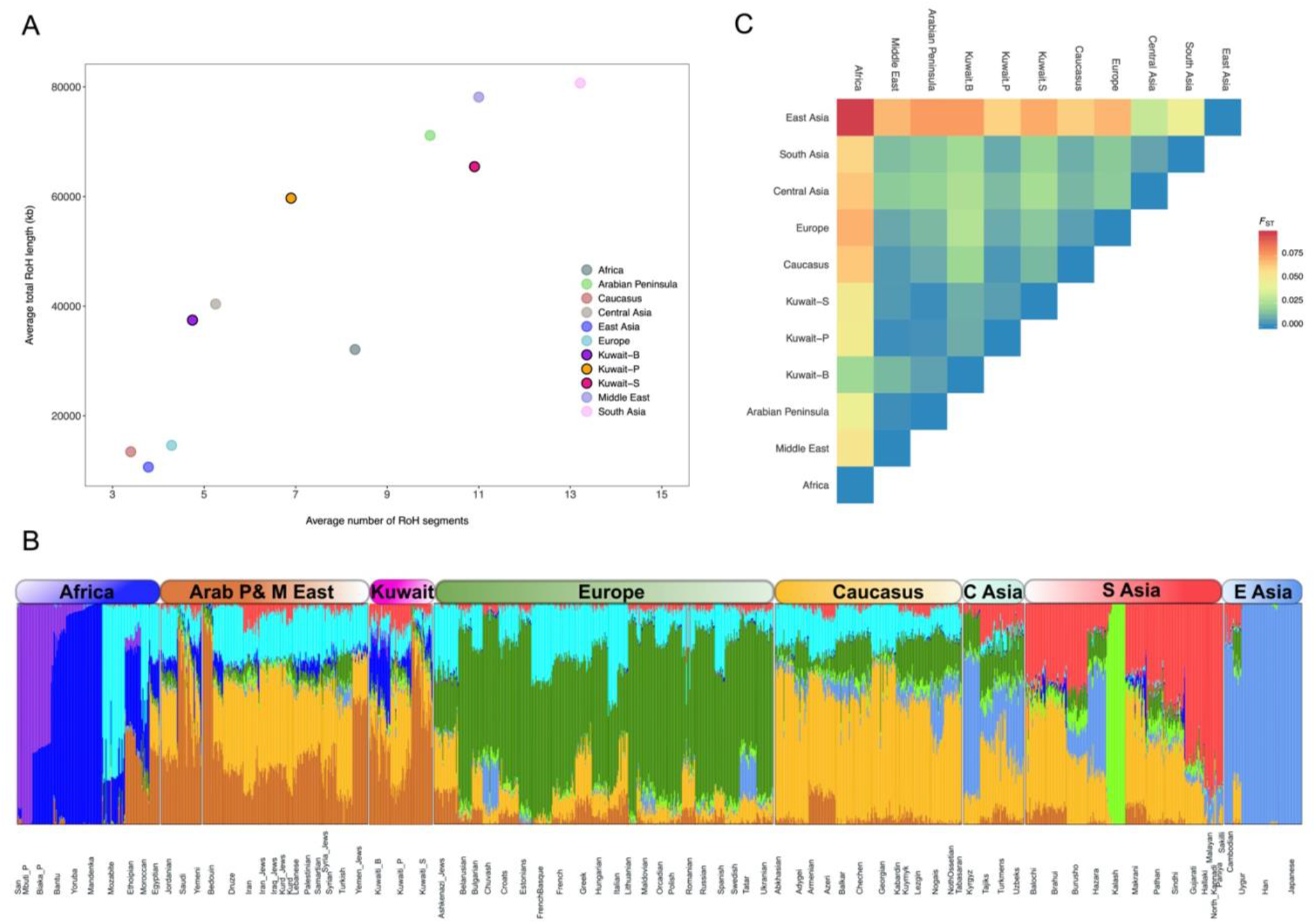
Population structure and ancestry components. **(A)** A scatter plot showing the average number of RoH segments and total RoH length in kb. **(B)** An ADMIXTURE plot of individual ancestry proportions at K = 9. **(C)** The mean pairwise *F*_ST_ values show the genetic distances between regional and continental populations.

### Datasets, merging and phasing

We merged our data with the published genome-wide SNP genotype data of global populations (Supplementary Fig. 1 and Supplementary Table 1) available from the Estonian Biocentre database (https://evolbio.ut.ee/). The combined dataset was filtered using PLINK version 1.9 [31] to include only autosomal SNPs with a minor allele frequency of >1%, a genotyping success rate of >97%, and only individuals with a genotyping success rate of >95%. The filtered combined dataset, which included 2139 individuals and 244,688 SNPs, was phased using the SHAPEIT algorithm [32] and used for further analyses. To avoid the effects of markers with a strong linkage disequilibrium, we thinned the marker set by removing SNPs with an *r*^2^ value of >0.4 using a sliding window of 200 SNPs, shifted at intervals of 25 SNPs. The pruned dataset yielded 155,744 SNPs that were used for the relevant population genetics analyses, including Wright’s *F*-statistic (*F*_ST_), principal component analysis (PCA), the ADMIXTURE tool for ancestry estimation, and runs of homozygosity (RoH).

**Table 1.** Admixture *f*_*3*_ (Source1, Source2; Target) to detect putative admixing sources in context of three assigned Kuwaiti population subgroups. “Source1” and “Source2” represent the admixing sources, and “Target” is any of the three Kuwaiti subgroups.

| Source1 | Source2 | Target | $f_3$ value | std err | Z |
| --- | --- | --- | --- | --- | --- |
| Yoruba | Saudi | Kuwait-S | -0.001966 | 0.000231 | -8.499 |
| Mandenka | Saudi | Kuwait-S | -0.001877 | 0.000224 | -8.368 |
| Ethiopian | Saudi | Kuwait-S | -0.000734 | 0.000124 | -5.905 |
| Ethiopian | Bedouin | Kuwait-B | -0.005084 | 0.00012 | -42.442 |
| Mandenka | Bedouin | Kuwait-B | -0.020919 | 0.000212 | -98.869 |
| Yoruba | Bedouin | Kuwait-B | -0.021847 | 0.00022 | -99.126 |
| Yoruba | Saudi | Kuwait-B | -0.023683 | 0.000262 | -90.526 |
| Ethiopian | Saudi | Kuwait-B | -0.005786 | 0.000131 | -44.068 |
| Mandenka | Saudi | Kuwait-B | -0.022782 | 0.000257 | -88.477 |
| Yoruba | Yemeni | Kuwait-B | -0.009428 | 0.000357 | -26.429 |
| Ethiopian | Yemeni | Kuwait-B | -0.002468 | 0.000146 | -16.902 |
| Yoruba | Kuwait-S | Kuwait-B | -0.02224 | 0.00019 | -117.171 |
| Yoruba | Kuwait-P | Kuwait-B | -0.02131 | 0.000198 | -107.865 |
| Mandenka | Kuwait-S | Kuwait-B | -0.021425 | 0.000186 | -115.478 |
| Mandenka | Kuwait-P | Kuwait-B | -0.020507 | 0.000192 | -106.721 |
| Ethiopian | Kuwait-S | Kuwait-B | -0.005541 | 0.000105 | -52.687 |
| Ethiopian | Kuwait-P | Kuwait-B | -0.006787 | 0.000105 | -64.91 |
| Romanian | Saudi | Kuwait-P | -0.000583 | 0.000098 | -5.943 |
| French | Saudi | Kuwait-P | -0.000487 | 0.000089 | -5.494 |
| Estonian | Saudi | Kuwait-P | -0.001725 | 0.000112 | -15.456 |
| Lezgin | Saudi | Kuwait-P | -0.002065 | 0.000087 | -23.815 |
| Sindhi | Saudi | Kuwait-P | -0.003833 | 0.00009 | -42.724 |
| Pathan | Saudi | Kuwait-P | -0.003741 | 0.00009 | -41.644 |
| Paniya | Saudi | Kuwait-P | -0.00385 | 0.000198 | -19.403 |
| Gujarati | Saudi | Kuwait-P | -0.004421 | 0.00011 | -40.124 |
| Iranian | Saudi | Kuwait-P | -0.000754 | 0.000074 | -10.228 |
| Orcadian | Saudi | Kuwait-P | -0.001071 | 0.000111 | -9.664 |
| Kurd | Saudi | Kuwait-P | -0.00108 | 0.000126 | -8.586 |
| Tajik | Saudi | Kuwait-P | -0.003027 | 0.000096 | -31.626 |
| Turkish | Saudi | Kuwait-P | -0.000703 | 0.000081 | -8.696 |
| French | Kuwait-S | Kuwait-P | -0.000923 | 0.000064 | -14.504 |
| Estonian | Kuwait-S | Kuwait-P | -0.0019 | 0.000077 | -24.625 |
| Armenian | Kuwait-S | Kuwait-P | -0.000629 | 0.000055 | -11.361 |
| Lezgin | Kuwait-S | Kuwait-P | -0.002432 | 0.000058 | -41.741 |
| Tajik | Kuwait-S | Kuwait-P | -0.00295 | 0.000065 | -45.271 |
| Gujarati | Kuwait-S | Kuwait-P | -0.003967 | 0.000075 | -53.191 |
| Paniya | Kuwait-S | Kuwait-P | -0.00316 | 0.000127 | -24.843 |
| Kurd | Kuwait-S | Kuwait-P | -0.001247 | 0.000088 | -14.18 |
| Iranian | Kuwait-S | Kuwait-P | -0.000938 | 0.000048 | -19.51 |
| Orcadian | Kuwait-S | Kuwait-P | -0.001474 | 0.000077 | -19.068 |

### Population structure analyses: *F*_*ST*_, PCA, ADMIXTURE and RoH

To explore the population genetic structure, we initially computed the mean pairwise *F*_*ST*_ differences between all population groups using the Weir and Cockerham method [33] implemented in PLINK version 1.9 [31]. Next, we conducted PCA of the linkage disequilibrium-pruned combined dataset using the smartpca program included with the EIGENSOFT software package version 6.1.4 [34,35]. Further, we ran an unsupervised structure-like analysis using the ADMIXTURE tool (version 1.3.0) [36] on the same dataset 25 times at different time intervals, with K values ranging from 2 to 12. Notably, K = 9 was the best supported K value as determined from the lowest cross-validation indexes. RoH was estimated using PLINK version 1.9 [31] with a sliding window of 100 SNPs (1000 kb), allowing for one heterozygous and five missing calls per window.

### Analyses to test admixture events and relative allele sharing: *f*_3_ and *f*_*4*_ *s*tatistics

We computed the *f*_3_ and *f*_*4*_ statistics using the *qp3Pop* and *qpDstat* programs (with *f*_*4*_ mode: YES) implemented in the ADMIXTOOLS software package [37]. A dataset containing 244,688 SNPs and 2139 individuals was used for analyses of the *f*-statistics of modern individuals. The dataset of modern genomes was merged with that of ancient specimens, which contained a combined total of 231,418 SNPs and 3697 individuals. To measure the genetic similarity of different Kuwaiti groups (i.e., modern and ancient), we computed the derived allele sharing of the Kuwaiti population using outgroup *f*_3_ in the form of *f*_3_(Mbuti; Pop1, X), where Pop1 is one of the three assigned groups of the Kuwaiti population, X is a modern or ancient West Eurasian population, and Mbuti is an outgroup (or Papuan to compare drift sharing between the Kuwait-B subgroup and other modern populations). Admixture *f*_3_ was also calculated to infer the plausible admixing sources in the history of the Kuwaiti population. We calculated the *f*_4_ statistic to evaluate the level of gene flow between contemporary populations of Kuwaitis and their regional neighbors and allele sharing between modern Kuwaitis and available published data of ancient individuals from surrounding regions. The *f*_*4*_ statistic was applied to numerous population combinations in the form of *f*_*4*_ (Pop1, Pop2, Pop3, Mbuti), where Pop1 and Pop2 are two Kuwaiti groups to be compared, while Pop3 is a modern or ancient population from a relevant region.

### Haplotype-based fine-scale analyses: ChromoPainter and fineSTRUCTURE

We used the phased dataset with 244,688 SNPs and 2139 individuals for haplotype-based analyses. The ChromoPainter tool, designed to identify haplotypes in sequence data, was used to “paint” each individual as a combination of all other sequences [38]. As a first step, we estimated the -n (recombination scaling constant) and -M (per site mutation rate) parameters by running the software’s EM option on a small subset of populations and five randomly selected chromosomes (3, 7, 10, 17, and 22), as described elsewhere [39]. The estimated values for the two parameters were -n = 510.86 and -M = 0.00033. Then, we executed the ChromoPainter tool in the “All vs All” mode (using the -a flag), where all individuals are considered as both donors and recipients.

Next, we analyzed the resulting painted dataset using the fineSTRUCTURE algorithm [38] to identify genetically homogenous clusters. We first ran the software performing 2 million burn-in iterations and 4 million MCMC iterations thinned every 10,000. This generated an MCMC file (.xml) that we used to build the tree structure using the option --T 1.

The analyzed individuals were initially classified into 233 clusters, which we reduced to increase the interpretability of subsequent analyses. More specifically, we iteratively “climbed the tree,” and the combined branches consisted of less than five clusters if at least one of them was composed of less than five individuals. The obtained tree was further refined by pooling together pairs or triplets of clusters if the pairwise total variation distance (TVD) based on the number of chunks shared among members of a branch was >0.035. After refinement, 40 clusters remained.

### Non-negative least square (NNLS)

Starting from the copying vectors obtained with the ChromoPainter tool, we reconstructed the ancestry profile of each cluster or individual by applying a slight modification of the NNLS function of R software version 3.5.1 (https://www.r-project.org/), as described elsewhere [39,40]. Therefore, for each individual belonging to a Kuwait cluster and for each of these clusters, we decomposed their ancestry as a mixture (with proportions summing to 1) of five (North/East Europe, Bedouins1, Yoruba, Druze, and North Africa clusters) and three (only North/East Europe, Bedouins1, and Yoruba) putative ancestral sources.

### Exploring the variability of Kuwaiti individuals via pairwise TVD analysis

To obtain a detailed picture of the variation underlying modern-day Kuwaiti population, we determined the pairwise TVD [40,41] among different individuals in specific clusters. TVD indicates the differences in ancestry profiles among individuals, where a high TVD value indicates high heterogeneity. First, we determined the TVD among individuals of the same cluster. Second, focusing only on Kuwaiti individuals, we determined the TVD among Kuwaitis and all the members of the respective clusters. Third, for each cluster, we determined the TVD only among Kuwaiti individuals. The analysis was performed in consideration of the number and length of genomic fragments inherited among individuals.

### Estimation of admixture dates

The times of admixture events were investigated using the GLOBETROTTER software [42]. We applied an “individual” approach by analyzing each Kuwaiti individual alone [43]. First, we estimated the time of the admixture event by applying the prop.ind=1, null.ind=0 approach to the 583 target individuals. Then, we performed 20 bootstrap iterations with the following settings: prop.ind=0, bootstrap.date.ind=1, and num.admixdates.bootstrap=1. For each of the inferred admixture events, we considered only those that were characterized by bootstrap values for the time of an admixture event between 1 and 400.

## Results

### Population structure

As populations inhabiting the Arabian Peninsula are well-known for their consanguinity, we compared the RoH patterns of the Kuwaiti population with those of regional and continental populations. Among the three Kuwaiti subgroups, the Kuwait-S subgroup was proximal to the Arabian Peninsula and the Middle East in terms of both the length and number of RoH segments (Fig. 1A), whereas the Kuwait-B and Kuwait-P subgroups were distant from the Kuwait-S subgroup, with the Kuwait-B subgroup displaying the lowest average length and number of RoH segments (Fig. 1A).

Furthermore, we applied the structure-like clustering algorithm ADMIXTURE to determine the detailed discrete genetic structure of the Kuwait population [36]. A given number of distinct ancestral populations is input as clusters (K) and the ADMIXTURE computes the genetic ancestry proportion of each individual for individual clusters or ancestral populations. We chose the best K value of 9 with the lowest cross-validation index. At K = 9, the Kuwaiti populations were characterized by six substantial ancestral components similar to their neighboring populations across the Arabian Peninsula and the Middle East (Fig. 1B). However, each of the three specified Kuwaiti subgroups harbor different proportions of these ancestral components. The brown (Arabian) component, which was shared by all populations of the Arabian Peninsula and the Middle East and maximized in Saudis and Bedouins, was the most prominent in Kuwait-S, followed by Kuwait-B and least in Kuwait-P subgroups. The Caucasus (orange), North African Mozabite-like (magenta), South Asian (red), and Kalash-like light green components were higher in the Kuwait-P subgroup (alike Iranians) than the two other Kuwaiti subgroups. The Kuwait-B subgroup was distinct among the Kuwaitis by having the highest overall African ancestry, which was dominated by the West African Mandenka/Yoruba-like blue component (Fig. 1B).

Assessment of mean pairwise *F*_ST_ differences reflected the regional affinity through the low degree of differentiation between the Kuwaiti subgroups and the Arabian Peninsula populations (Fig. 1C). The Kuwait-P subgroup had a shorter genetic distance (*F*_ST_ < 0.01) from Arabians, Middle Easterners, Caucasians, South Asians, and Europeans, and the Kuwait-B subgroup with Arabians, and the Kuwait-S subgroup with Arabians and Middle Easterners. The highest degree of differentiation (*F*_ST_ > 0.05) was observed between the Kuwait-P/Kuwait-S subgroups and Africans/East Asians. The Kuwait-B subgroup was genetically distant from East Asians, but not Africans (Fig. 1C). The individual population-wise *F*_ST_ differences are presented in Supplementary Fig. 2.

**Fig. 2.**
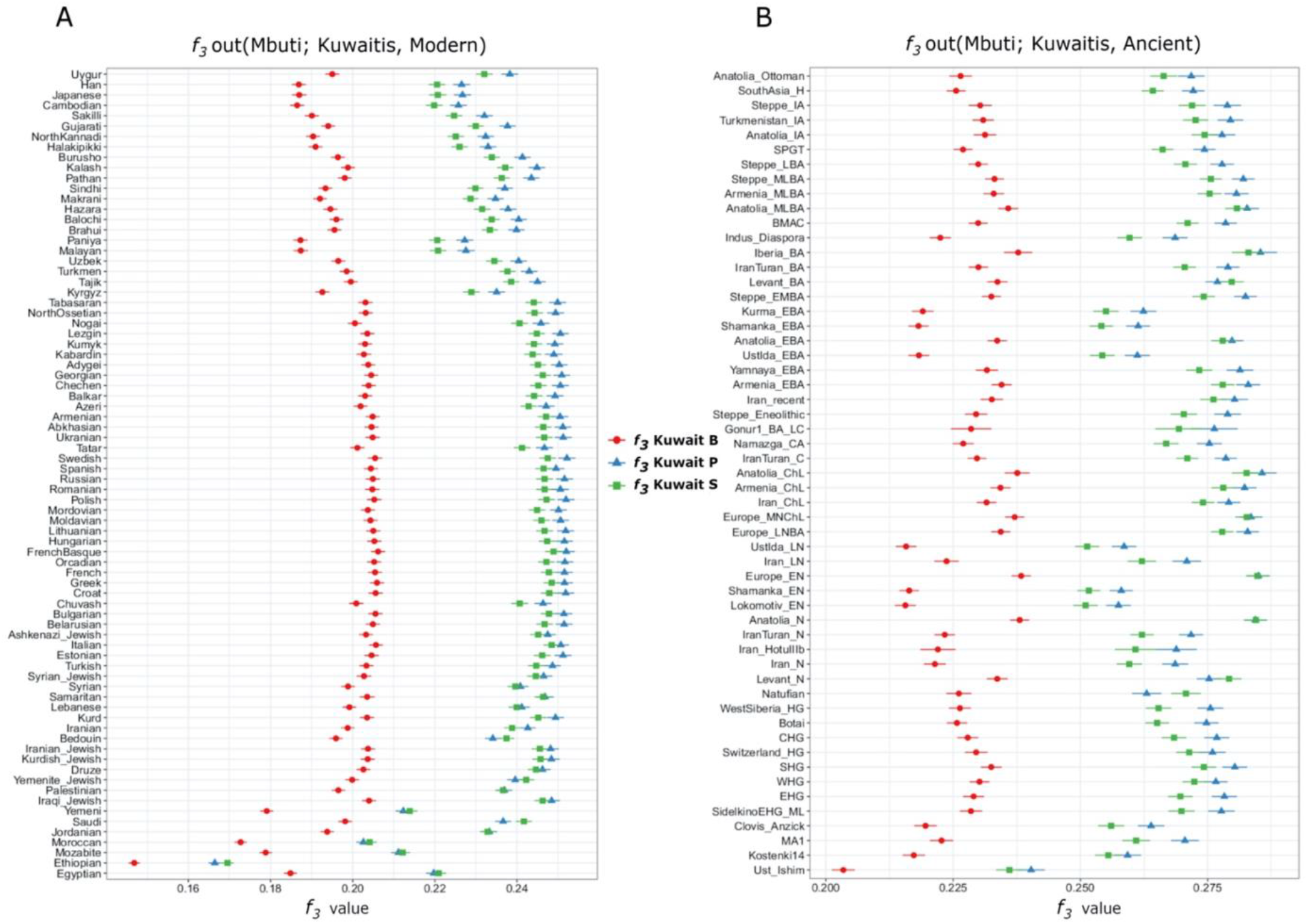
Outgroup *f*_*3*_ (Mbuti; Pop1, X) results of the three subgroups of Kuwaiti populations with Mbuti as an outgroup. **(A)** Comparison of the shared drift of the Kuwaiti populations to that of other modern individuals. **(B)** Relative affinities to ancient individuals from West Eurasia.

PCA was performed to evaluate the relationships among global and regional populations, which inferred the geographical affinity of all three Kuwaiti subgroups from their distributions in the close vicinity of their Arabian and Middle Eastern neighbors (Supplementary Fig. 3). As the allele frequency-based PCA did not reveal the heterogeneous population structure at the finest level, we further performed PCA of the haplotype copying vectors (Fig. 3) together with other haplotype-based investigations.

**Fig. 3.**
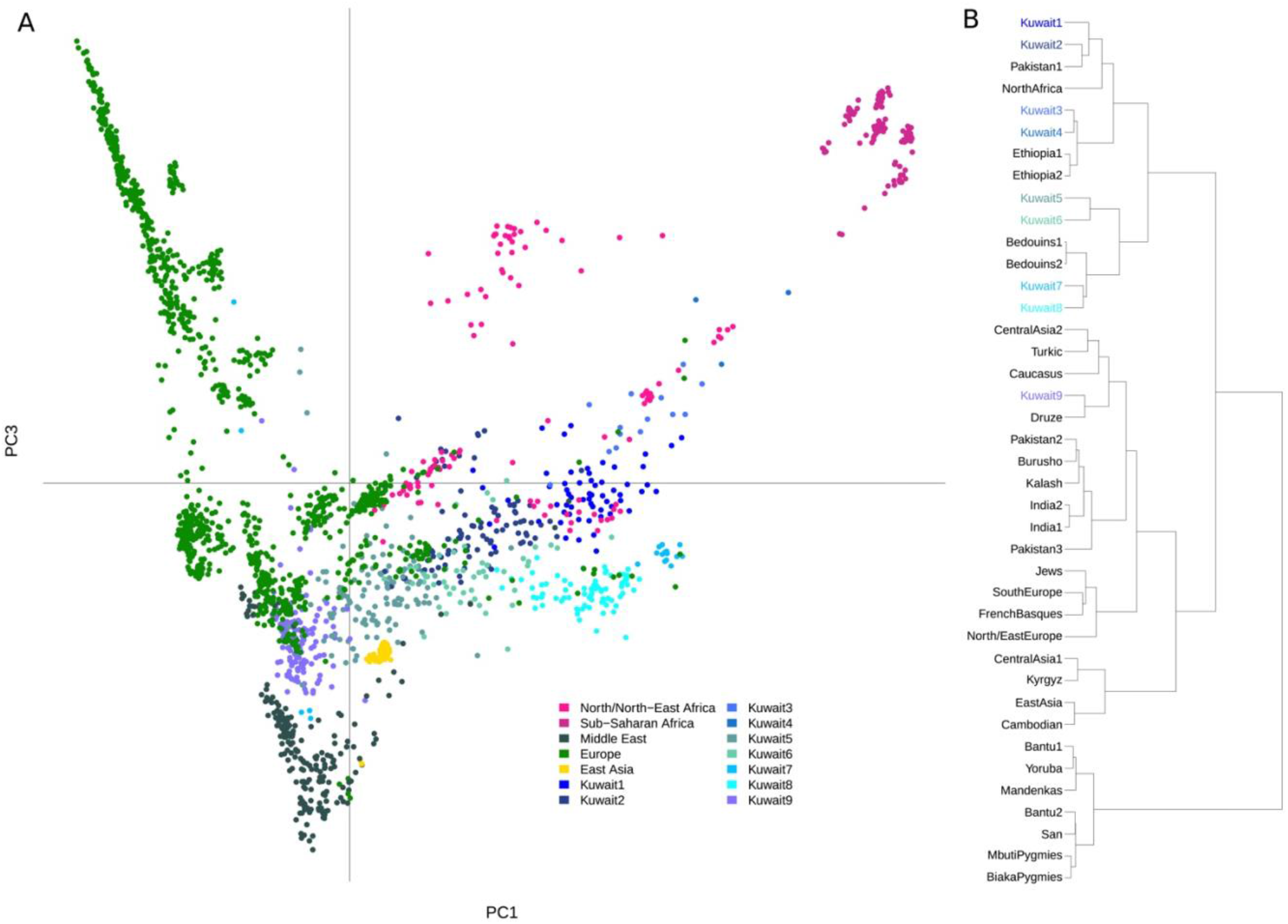
**(A)** PCA based on haplotype sharing. The chunkcount coancestry matrix obtained with the ChromoPainter was used to perform PCA analysis with *prcomp* in R. The first and third components are shown in the scatterplot. **(B)** The refined fineSTRUCTURE tree consisted of 40 homogeneous clusters. Clusters containing more than two Kuwaiti individuals are presented in various colors, as shown in the legend.

### Drift and allelic sharing of the Kuwaiti population with modern and ancient individuals

The *f*_*3*_ admixture results provide distinct patterns of the histories of all three Kuwaiti subgroups, wherein the Kuwait-B subgroup had a significant admixture signal (i.e., *Z*-score < -3) when one of the ancestral sources was either Bedouin or Saudi, while the second source was a different African population (Table 1). Both the Kuwait-S and Kuwait-P subgroups had a common ancestral source in Saudis, but not Bedouins, while the second source varied. The signal for the Kuwait-S subgroup was significant only when the second ancestral source was African, whereas the Kuwait-P subgroup had a significant admixture signal when the second ancestral source was a population from South Asia, Parsi, Caucasus, or Central Asia. Hence, among the three subgroups, the Kuwait-S is probably related to an ancestral basal population group of the Arabian Peninsula.

By *f*_*3*_ outgroup analysis with modern populations using Mbuti as an outgroup, we observed that all three Kuwaiti subgroups had an identical declining pattern of shared drift from Europe to East Asia (Fig. 2A). The results also showed that the Kuwait-B subgroup had a distinctly lower shared drift than both the Kuwait-P and Kuwait-S subgroups, in regard to all modern populations across the globe. Compared with the Kuwait-S subgroup, the Kuwait-P subgroup had a higher shared drift with most Eurasian populations, with the exception of those from the Arabian Peninsula and Africa. However, the Kuwait-S subgroup had a higher shared drift with populations with deeper Arabian and African genetic backgrounds. Interestingly, among the three subgroups, the Kuwait-B subgroup had the least drift sharing with Africans, which can probably be explained by the higher masking of African-related alleles from the outgroup. To verify this hypothesis, we performed another *f*_*3*_ analysis (Supplementary Fig. 4) using Papuans (instead of Mbuti) as an outgroup underlying the idea of an early split of Papuans from other non-Africans. Consequently, we observed that the drift sharing was greater for the Kuwait-B subgroup with Africans than the other two Kuwaiti groups. This drift sharing of the Kuwait-B subgroup with Africans was relatively higher with Yoruba/Mandenka/Bantu/African Pygmy populations than that with Ethiopian/Moroccan/Mozabite populations.

**Fig. 4.**
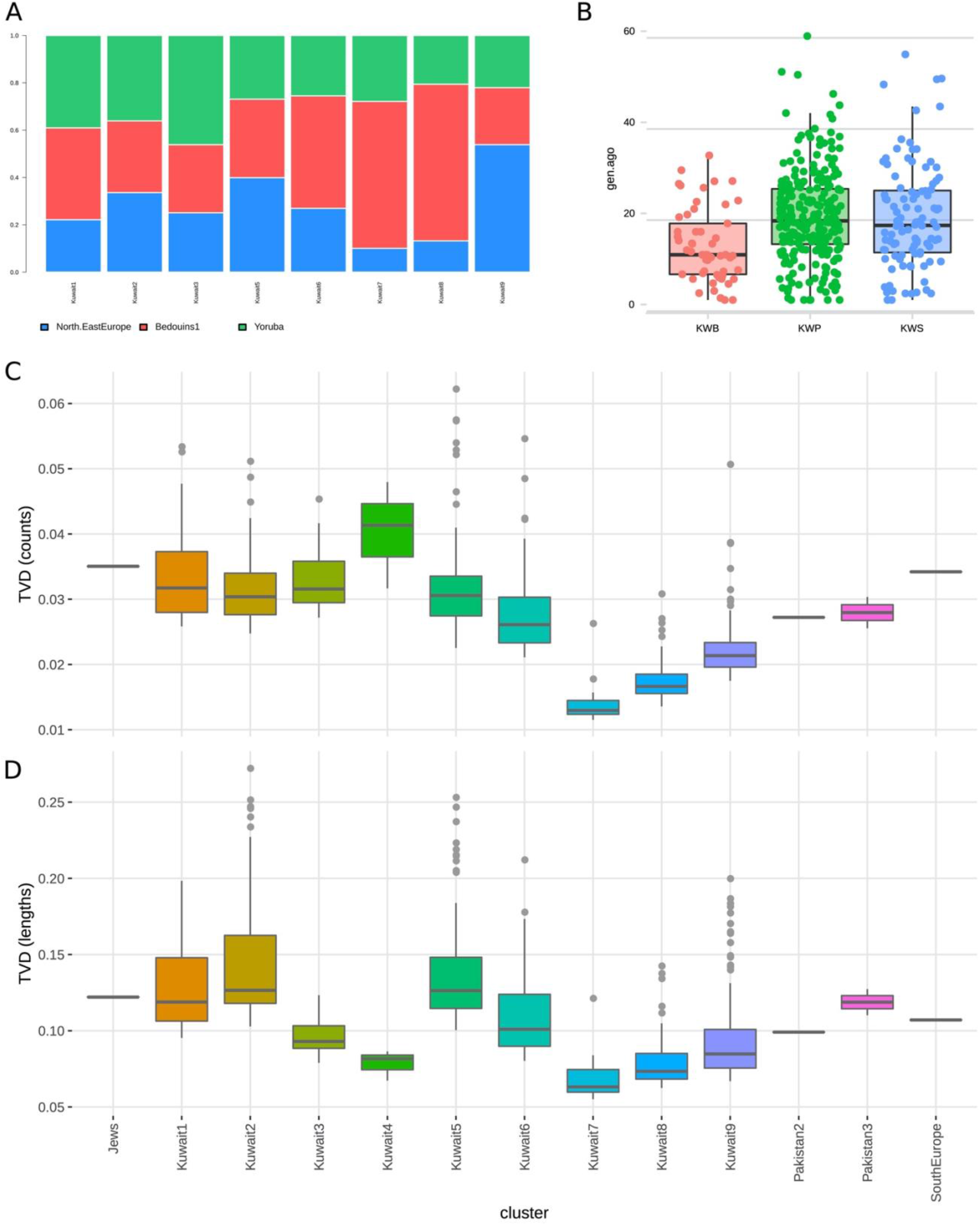
**(A)** Ancestry proportions of the main eight Kuwaiti clusters as inferred by NNLS analysis using the North/East Europe, Bedouins, and Yoruba clusters as putative sources. **(B)** Inferred admixture dates by GLOBETROTTER population-based analysis. **(C & D)** Intra-cluster TVD of the number and length of genomic fragments of Kuwaitis and individuals from the same cluster.

In the context of the *f*_*3*_ outgroup analysis with ancient West Eurasian individuals, the Kuwait-B subgroup again stood apart from the other subgroups due to the low derived allele sharing with all analyzed ancient genomes (Fig. 2B). The Kuwait-P subgroup had the highest genetic affinity with Iberian_BA or Anatolian_ChL individuals but showed higher genetic relatedness with most of the ancient individuals compared with the Kuwait-B and Kuwait-S subgroups. With respect to the Europe_EN and Europe_MnChl, the Kuwait-P and Kuwait-S subgroups shared equal amounts of derived alleles. Interestingly, the Kuwait-S subgroup had higher genetic sharing with the Levant_N, Levant_BA, and Natufians than the Kuwait-P and Kuwait-B subgroups. In general, all three subgroups showed a pattern of higher genetic affinity with ancient individuals from the Steppe and Caucasus regions.

The *f*_*4*_ statistical results of the modern genomes (Supplementary Table 2) showed higher allelic sharing (gene flow) of the Kuwait-P and Kuwait-S subgroups with all Eurasian populations, compared with the Kuwait-B subgroup. Kuwait-P was found to have the highest sharing of alleles with all Eurasian populations, with the exception of those from the Arabian Peninsula and Africa. Populations of the Arabian Peninsula and Africa mostly shared higher alleles with the Kuwait-S subgroup than with the Kuwait-P and Kuwait-B subgroups. Given the pronounced African genetic component of the Kuwait-B subgroup in the ADMIXTURE results, the less allelic sharing in *f*-stats could be the consequence of higher masking of African-like alleles.

The *f*_*4*_ statistical analysis with aDNA also disclosed a similar pattern, as the Kuwait-P and Kuwait-S subgroups shared higher alleles with all ancient individuals than with the Kuwait-B subgroup (Supplementary Table 3). Kuwait-P shared more alleles with almost all ancient individuals than the Kuwait-S subgroup, except in the context of the Levant_N, Levant_BA, and Natufian, which shared more alleles with the Kuwait-S subgroup than with the Kuwait-P subgroup. However, with regard to the Europe_EN and Anatolia_N, both the Kuwait-P and Kuwait-S subgroups shared almost equivalent numbers of alleles.

### Distinct genetic heterogeneity in the Kuwaiti population

To understand the fine-scale population structure and recent admixture history, we scrutinized the haplotype-sharing patterns among Kuwaiti individuals, which were pooled into three subgroups, as in our previous investigations. Through fineSTRUCTURE analysis, we inferred the existence of eight different groups that included at least five Kuwaiti individuals (Fig. 3B). Of these, three (Kuwait1-3) were part of a group of clusters consisted of all individuals from the Kuwait-B subgroup including three that formed a minor group. Four additional (Kuwait4-7) groups comprised a group of clusters that included groups from the Arabian Peninsula, which included 38% individuals of the Kuwait-P subgroup and 75% of the Kuwait-S subgroup. Moreover, 33% of the Kuwait-P subgroup were part of a larger group that included Jews and populations from Anatolia, the Caucasus, and the Levant, which formed a sister group with the Druze population. This cluster scheme confirmed that the previous clustering postulating the existence of three different groups had only partially captured the high genetic variation of Kuwaitis.

The relationships among populations were further explored by PCA based on the chunkcount coancestry matrix. As shown in the PCA scatter plot presented in Fig. 3A, PC1 and PC3 accounted for 0.24% and 0.08% of the variation, respectively. In general, the fineSTRUCTRE clustering of Kuwaiti individuals mirrors the assemblies of individuals of different Kuwaiti groups with corresponding populations of the same fineSTRUCTURE sub-cluster, rather than with each other.

The allele frequency and haplotype-based results further divided the genetic structure of the Kuwaiti population, highlighting the high genetic variability among the three Kuwaiti subgroups. Therefore, to ensure that the three subgroups sufficiently accounted for the existent genetic variability of the Kuwaiti population, we explored the pairwise TVD distribution at different levels. Basically, we calculated the TVD between individuals of a cluster considering haplotypic copying vectors of the individuals. When the intra-cluster TVD was taken into consideration, most of the Kuwaiti samples had the highest TVD among all clusters in the dataset, revealing strong heterogeneity (Supplementary Fig. 5). This heterogeneity was still evident when only the TVD (a) among Kuwaitis and all the members of the same clusters (Fig. 4C and 4D) and (b) among Kuwaitis of the same cluster (Supplementary Fig. 6) were explored, implying that the high diversity of the clusters with large numbers of Kuwaitis is, at least in part, driven by the high heterogeneity among native Kuwaitis, rather than by differences among individuals from the Arabian Peninsula.

### Ancestral genetic components of the Kuwaiti population clusters

The extreme variation of the analyzed samples was also confirmed when the ancestry of the eight clusters was explored with the NNLS function (Fig. 4A). Namely, the proportion of inferred European, African, and Middle East/Arabian was quite variable. More specifically, the Kuwait1, Kuwait2, and Kuwait3 clusters were characterized by a similar proportion of the three ancestries, with a slightly higher contribution from Africa (39%, 36%, and 46% from the Yoruba cluster, respectively). In contrast, clusters Kuwait6, Kuwait7, and Kuwait8 were characterized by a large proportion of Middle East/Arabian ancestry (47%, 62%, and 66% from the Bedouins1 cluster, respectively). The Kuwait5 cluster had similar contributions from all sources, with the greatest contribution from Europe (26%, 33% and 39% from the Yoruba, Bedouins1, and North/East Europe clusters, respectively). Finally, the Kuwait9 cluster had the highest European ancestry (54% from the North/East Europe cluster). The genetic contributions of the Druze and North Africa clusters were also evaluated, but the proportions did not vary much among the eight Kuwaiti clusters (Supplementary Fig. 7).

### Recent admixture events

To identify the admixture events with different sources for the three Kuwaiti subgroups, we analyzed their haplotype data using the GLOBETROTTER software [42], which disclosed different genetic profiles of the major and minor ancestral sources of the three subgroups (Supplementary Fig. 8). The major ancestral sources of the Kuwait-P subgroup were Kuwait, Europe, and South Asia, those of the Kuwait-B subgroup were Africa and Kuwait, and that of the Kuwait-S subgroup was Kuwait. We found different minor ancestral sources of Africa, such as Mandenka and some hunter-gatherers, for the Kuwait-S subgroup, which also had high Central Asian ancestral sources (Supplementary Fig. 8A). On the other hand, the Kuwait-B subgroup underwent several African admixture events (Supplementary Fig. 8B). Moreover, while inferring the time of admixture events of the three groups, we observed the GLOBETROTTER results showing significantly more recent admixture events for the Kuwait-B subgroup than for the Kuwait-P and Kuwait-S subgroups (Fig. 4B, Wilcoxon test with Bonferroni-adjusted *p*-values: KWB vs. KWP = 0.000012; KWB vs. KWS = 0.0012). More specifically, the mean inferred time for the Kuwait-B subgroup was 13 generations ago (5%-95%: 2-27 generations ago), and similar values were observed for the Kuwait-P and Kuwait-S subgroups (∼19 generations; 5%-95%: 4-37 and 3-43 generations ago, respectively) (Fig. 4B).

## Discussion

The results of the classical allele frequency-based analyses conducted in the present study are in agreement with our previous genome-wide study of the genetic substructure of the Kuwaiti population, which showed genetic heterogeneity [24]. Additionally, the RoH were relatively long and frequent in the Kuwaiti population, but also divergent among the subgroups, which is rather intriguing considering the high prevalence of consanguinity in the studied population. Thus, we applied refined statistical approaches to analyze the genetic structure of the Kuwaiti population in the context of available ancient samples and haplotypes of modern genomes. As the “within” population structure is embedded within haplotypes, we resolved the extent of genetic heterogeneity by fine-scale, high-resolution, haplotype-based analyses. We applied the ChromoPainter tool and fineSTRUCTURE algorithm to uncover the hidden genetic structure “within” the Kuwaiti population.

These methods exploit the rich information available within haplotypes to identify clusters of genetically distinct individuals with a resolution that cannot be attained with the use of allele frequency-based methods. Through this approach, we could identify distinct genetic clusters of individuals that strongly segregate within the three Kuwaiti subgroups.

The results of *f*_*3*_ and *f*_*4*_ confirmed the existence of a heterogeneous genetic pattern among the Kuwaitis, signaling out a probable impact of distinct population dynamics that characterized the current genetic diversities of different Kuwaiti subgroups. The *f*_*3*_ admixture results (Table 1) showed a distinct set of ancestral sources for the Kuwait-B subgroup, rather than for the Kuwait-P and Kuwait-S subgroups, suggesting a different genetic background of the Kuwait-B subgroup related to ancestors of contemporary Bedouins and Africans. Notably, a similar higher African-related genetic background of the Kuwait-B subgroup was also observed in the results of ADMIXTURE and CP-NNLS analyses. However, these results did not replicate similar patterns in the outgroup *f*_*3*_ and *f*_*4*_ statistics with African Mbuti as an outgroup, probably due to the masking of alleles shared with the outgroup. This higher masking of alleles related to the outgroup in the Kuwait-B subgroup plausibly reflects a greater number of African-related alleles among individuals in the Kuwait-B subgroup possibly due to a recent admixture event. There was greater genetic affinity between the Kuwait-P and Kuwait-S subgroups. With regard to other modern and ancient populations (Fig. 3A and 3B), the significant admixture signals with the use of the Saudi population as an admixture source (Table 1) indicate a relatively higher common genetic background of these two groups compared with the Kuwait-B subgroup. However, considering the relatively recent origin of the Kuwaiti population from the Saudi people, the discrepancies in the admixture signal, shared drift, and allele sharing pattern of the Kuwait-B subgroup, particularly in the context of both modern and ancient individuals, can be interpreted as a consequence of later gene flow from African-related populations to the Kuwait-B subgroup. This is obvious from the lowest number and length of RoH segments in the Kuwait-B subgroup (Fig. 1A), suggesting higher interactions with outsiders.

Meanwhile, the visibly higher genetic affinity of the Kuwait-S subgroup to the Arabian Peninsula and African populations, compared with the Kuwait-P subgroup in admixture *f*_*3*_ (Table 1), the shared drift (Fig. 2A) and allele sharing (Supplementary Table 2) patterns suggest a varied population history for both and a lesser extent of intermixing. In fact, the highest amount of RoH segments (both in the average number and length) of the Kuwait-S subgroup is also in agreement with inbreeding and negligible interactions with other groups. Such consanguinity is plausibly a causal factor among individuals in the Kuwait-S subgroup of being much closer to their ancestral source groups from the Arabian Peninsula, which is also supported by the admixture *f*_*3*_ results showing that among the three subgroups, the Kuwait-S subgroup was more of a basal group. The greater affinity of the Kuwait-S subgroup with populations from the Arabian Peninsula is also corroborated by the *f*_*3*_ outgroup results with modern data and even with ancient individuals where the Kuwait-S subgroup had greater affinity to ancient Levant farmers and hunter-gatherers than the Kuwait-P subgroup (Fig. 2B). Meanwhile, the genetic affinity of the Kuwait-P subgroup to European and Caucasus populations is in agreement with the Persian-related genetic background. Considering this fact, it is anticipated that the Kuwait-P subgroup would be more closely related to ancient populations from the Steppe, Caucasus, and Iran, with lesser genetic affinity to ancient Levantines and Natufian populations than the Kuwait-S subgroup, as reflected in outgroup *f*_*3*_ with ancient specimens. It was intriguing to find less allele sharing of the Kuwait-B subgroup with Africans in *f*_*3*_ with Mbuti as outgroup, especially considering the higher African genetic components as determined by ADMIXTURE analysis. However, in the additional *f*_*3*_ using Papuans as the outgroup (Supplementary Fig. 4), strikingly, the Kuwait-B subgroup had the highest degree of drift sharing with West Africans, suggesting a recent gene flow between these groups.

Taking into account that both the West and East African populations were transported to the Middle East, Arabia, and Indian Ocean during the slave trade in the 15^th^-19^th^ centuries [44,45], the *f*_*3*_ admixture results of the Kuwait-B and Kuwait-S subgroups verified the impact of the slave trade on populations inhabiting the Arabian Peninsula. Moreover, considering the NNLS results of higher ancestry contribution from Yoruba in Kuwait1, 2, and 3 clusters (with most Kuwait-B individuals), and the low level of variation in North African genetic ancestry profile for all eight Kuwaiti clusters in addition to the admixture signal with Mandenka, we infer a possible recent admixture event in the Kuwait-B subgroup with a West African (Mandenka or Yoruba) or sub-Saharan African group.

Furthermore, the GLOBETROTTER results reinforce different major and minor ancestral sources of all three Kuwaiti subgroups (Supplementary Fig. 8A) and varied admixture events (Supplementary Fig. 8B). The higher African source profile of the Kuwait-B subgroup suggests a more recent admixture event within the last ∼10 generations (∼300 years) for the Kuwait-B subgroup than the other two subgroups (Fig. 4B). The Kuwait-P and Kuwait-S subgroups share a similar single date of admixture approximately 18 generations (∼500 years) ago (Fig. 4B; Supplementary Fig. 8B). These admixture events signify the role of the slave trade and Arabian maritime dominance [45-47] by the genetic footprint in present-day Kuwaitis as was expected, but it is almost obvious that the Kuwait-B subgroup is probably one of the most exogamous groups inhabiting the Arabian Peninsula with a different and more recent history of intermixing that plausibly involved slaves from Western or sub-Saharan Africa [47]. Moreover, the fineSTRUCTURE and PCA identified distinct genetic clusters of individuals that strongly segregated within the three Kuwaiti subgroups, indicating a much deeper and distinct level of genetic heterogeneity among present-day Kuwaitis.

The TVD results of the differences in genetic ancestry profiles of individuals of a cluster provided a better picture of the genetic heterogeneity of the Kuwaiti population. Notably, the intra-cluster TVD value was the highest (both in length and number) for Kuwaiti populations inhabiting the Arabian Peninsula and remained highest when the TVD was calculated only among Kuwaiti individuals in a single cluster. Consequently, the results indicate that the heterogeneity of clusters was due to the Kuwaiti population, suggesting that Kuwaitis are one of the most genetically heterogeneous populations inhabiting the Arabian Peninsula. NNLS analysis also confirmed the genetic heterogeneity of the Kuwaiti population by showing the differential amount of genetic ancestries that contributed to each of these clusters by three different major donor populations of Yoruba, Bedouins, and North-East Europeans.

In general, Kuwaiti populations are demographically characterized by large families and a remarkable rate of consanguinity, which is a potential threat to human health, especially in the context of rare autosomal recessive genetic diseases. As there have been historical population migrations in this region, a complex genetic diversity is expected, which was clearly reflected in this study. Our thorough investigations on the Kuwaiti population genetic structure at the finest scale highlights the precise genetic history and distinct heterogeneity of the Kuwaiti people, which could enormously aid in the systematic discovery of population- and/or family-specific diseases, especially in deciphering deleterious founder genetic variations. Overall, our study presents the fine-scale genetic structure of the distinctively heterogeneous Kuwaiti population and further highlights the recent historical population influx and gene flow from Western/sub-Saharan Africa to the Arabian Peninsula region.

## Acknowledgments

This work was supported by a research grant to the Dasman Diabetes Institute from the Kuwait Foundation for the Advancement of Sciences (grant no. RA 2015-022). We thank the members of the National Dasman Diabetes BioBank Core Facility for sample processing and DNA extraction. We acknowledge Fadi Alkayal and Motasem Melhem for genotyping assays. We thank Prof. Gyaneshwer Chaubey for his support during the initial stages of this work. We thank Dr. Luca Pagani for his useful comments and suggestions.

## Author Contributions

TAT and ME designed the study. OA was responsible for participant recruitment, sample collection and genotyping. PH processed the raw genotype data. OA and PH were involved in subgroup classification of the samples. ME, AKP, LO, and FM analyzed the data. ME, AKP, and MM contributed to the interpretation of results. ME and AKP wrote the main article. TAT, FAM, LO, and FM contributed to the writing of the article. FAM provided the required resources and critically reviewed and approved the final version of the article. All authors reviewed and approved the final version of the article prior to submission.

