## Supplemental Information for "Fine-scale population structure reveals high genetic heterogeneity of the Kuwaiti population in the Arabian Peninsula"

### Supplementary Information

#### Supplementary Figures

**Supplementary Fig. 1.** (A) Geographical location of the populations included in the study. (B) Sample information, quality control measures and the population genetics analyses performed in the study.

**Supplementary Fig. 2.** The mean pairwise $F_{ST}$ values showing genetic distance between individual populations.

**Supplementary Fig. 3.** Principal Component Analysis plot representing the first two principal components, PC1 and PC2, accounting for 2.11% and 1.72% of the variation respectively.

**Supplementary Fig. 4.** $f_3$ -statistics with Papuans as outgroup, comparing the genetic affinity of the Kuwaiti population subgroups to other modern populations.

**Supplementary Fig. 5.** Intra cluster Total Variation Distance for number and length of genomic fragments among the 40 clusters inferred by fineSTRUCTURE. (A) TVD estimated on number of fragments. (B) TVD estimated on length of fragments.

**Supplementary Fig. 6.** Intra cluster Total Variation Distance for number and length of genomic fragments among Kuwaiti individuals from the same cluster. Lower boxplots refer to TVD estimated on number of fragments and the upper boxplots refer to TVD estimated on length of fragments.

**Supplementary Fig. 7.** Ancestry proportions of the main eight Kuwait clusters as inferred by NNLS analysis using North/East Europe, Bedouins, Yoruba, Druze and North Africa clusters as putative sources.

**Supplementary Fig. 8.** Inferred admixture sources and times by GLOBETROTTER population based analysis. (A) The x and y axes show the minor and major sources, respectively, as inferred by GLOBETROTTER. (B) Type of admixture.

#### Supplementary Tables

**Supplementary Table 1.** Details of the populations included in the study.

**Supplementary Table 2.** $f_4$-statistics comparing the genetic affinity of the Kuwait population subgroups to other modern populations.

**Supplementary Table 3.** $f_4$-statistics for relative allele sharing of the Kuwait population subgroups to ancient West Eurasian individuals.

**Supplementary Fig. 1.** (A) Geographical location of the populations included in the study. (B) Sample information, quality control measures and the population genetics analyses performed in the study.

A

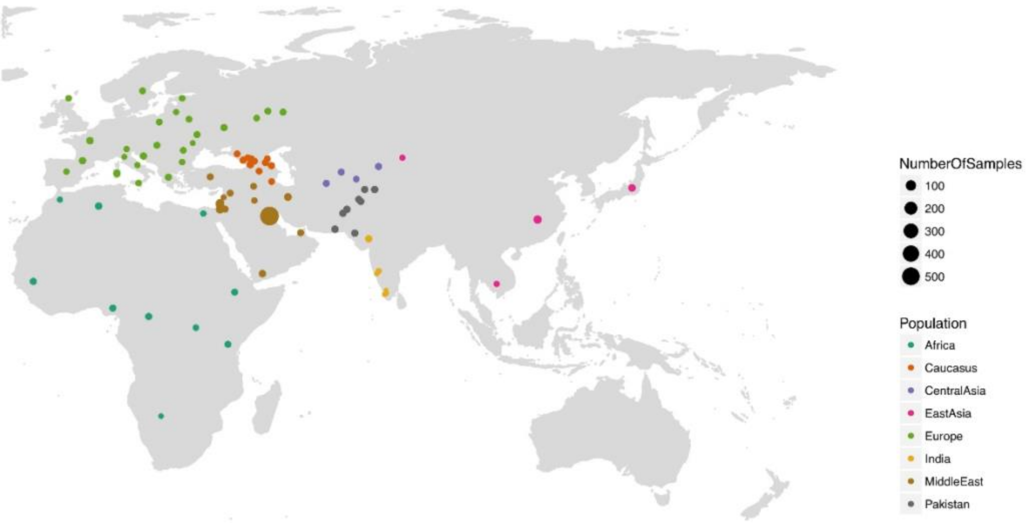


B

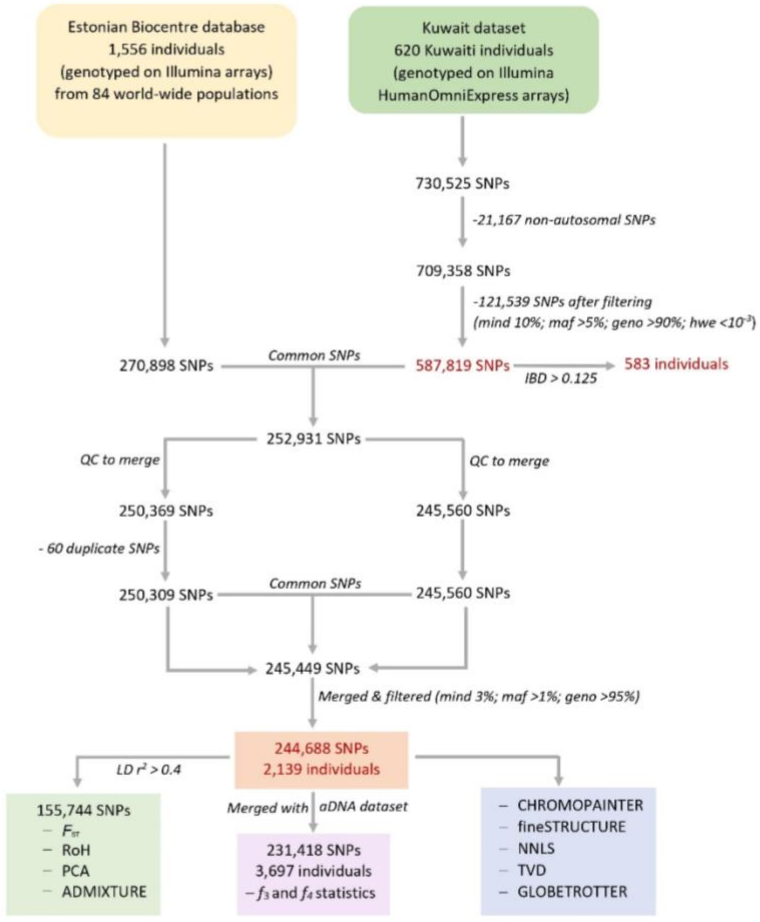

**Supplementary Fig. 2.** The mean pairwise $F_{ST}$ values showing genetic distance between individual populations.

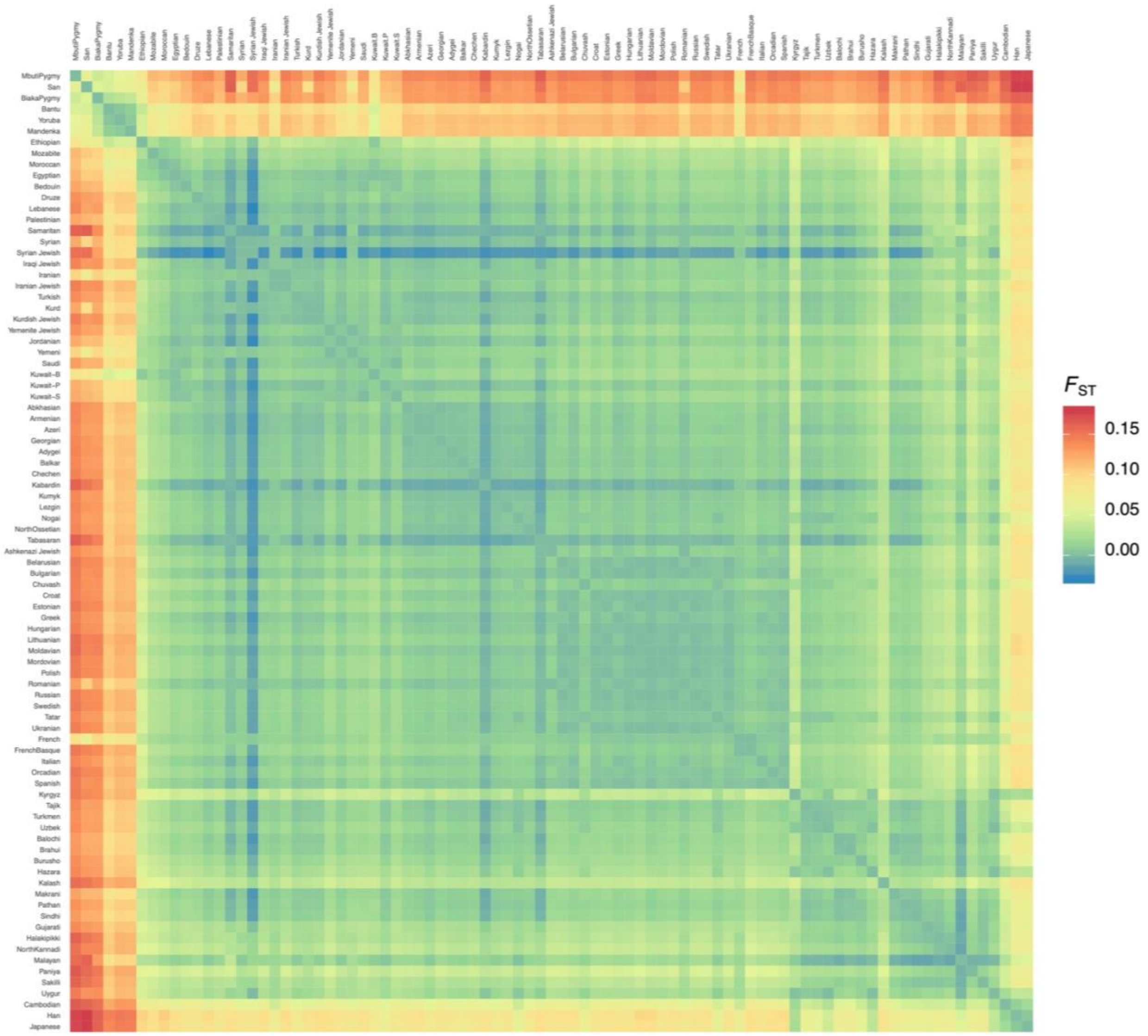

**Supplementary Fig. 3.** Principal Component Analysis plot based on allele frequency representing the first two principal components, PC1 and PC2, accounting for 2.11% and 1.72% of the variation respectively.

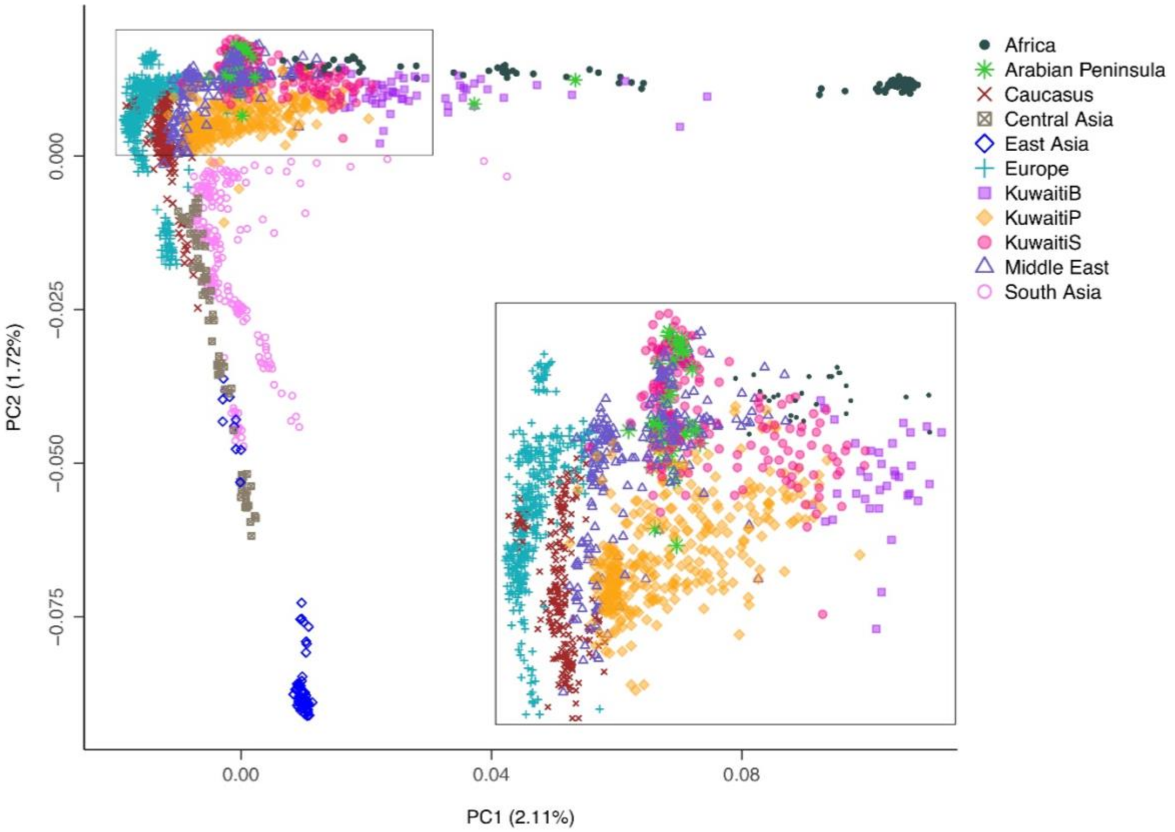

**Supplementary Fig. 4.** *f*3 -statistics with Papuans as outgroup in the form D(Papauans; Pop1, X) where Pop1 represents a Kuwaiti subgroup and X is a present-day population, comparing the genetic affinity of the Kuwait population subgroups to other global modern populations.

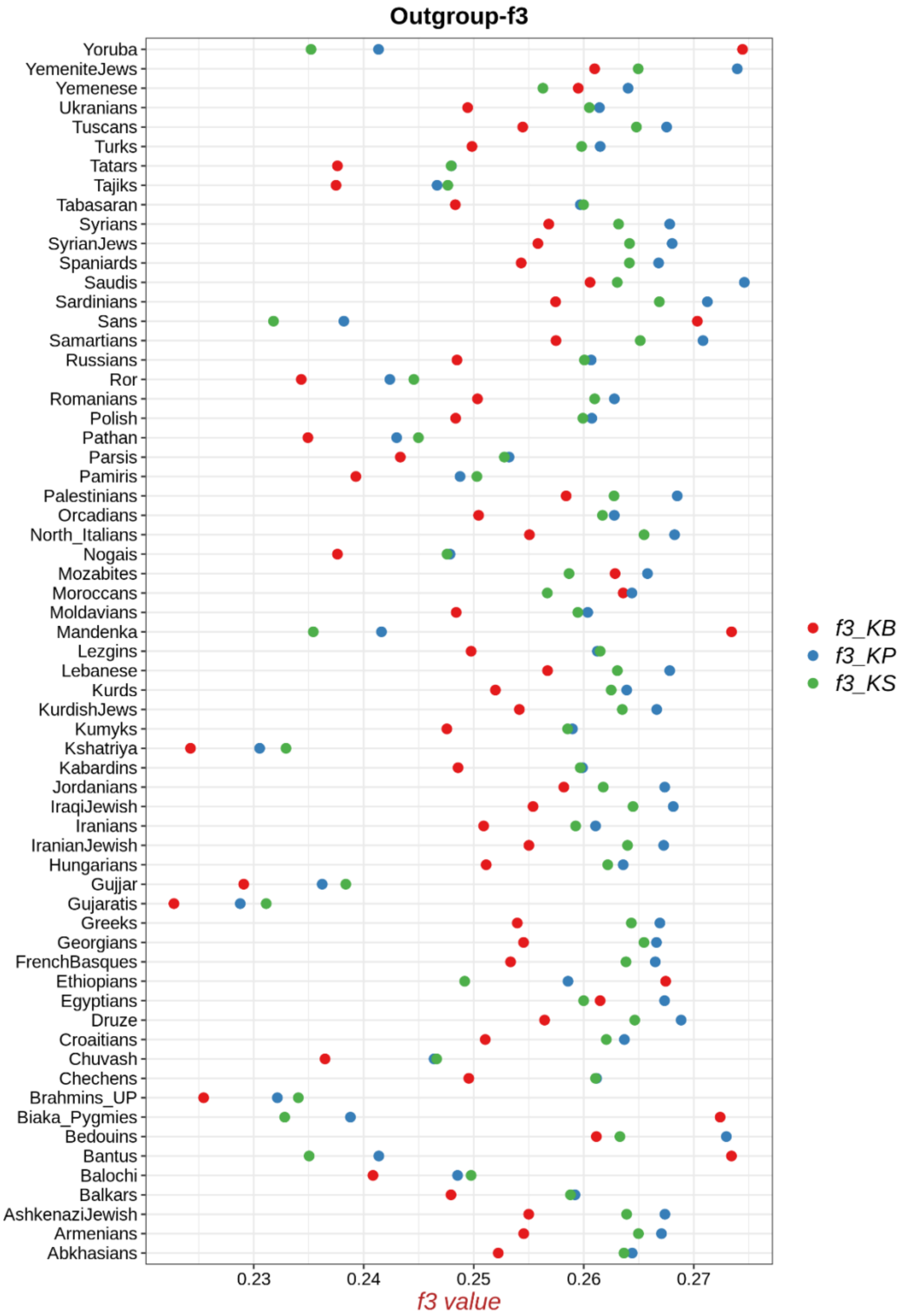

**Supplementary Fig. 5.** Intra cluster Total Variation Distance (TVD) for number and length of genomic fragments among the 40 clusters inferred by fineSTRUCTURE. (A) TVD estimated on number of fragments. (B) TVD estimated on length of fragments.

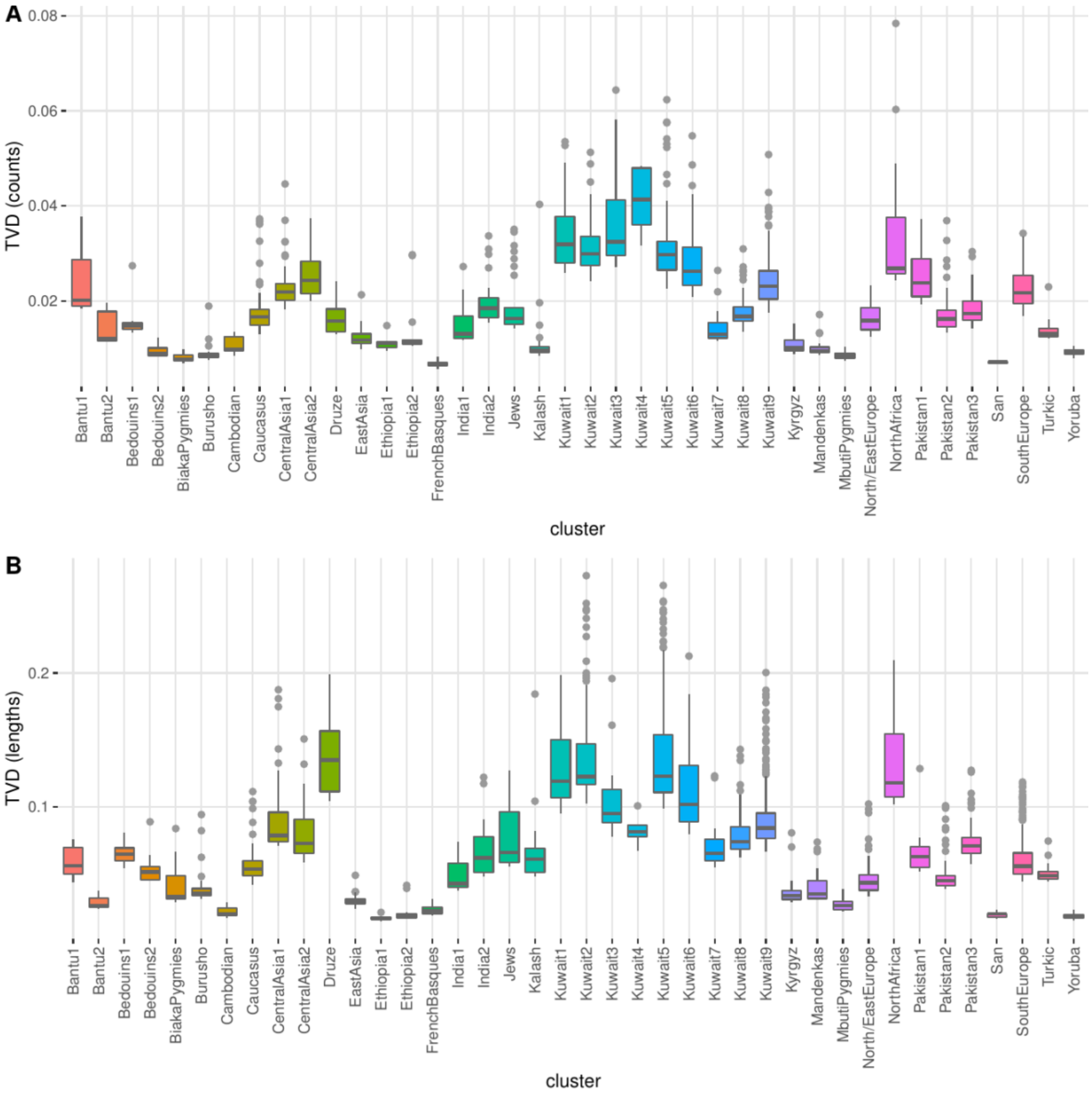

**Supplementary Fig. 6.** Intra cluster Total Variation Distance for number and length of genomic fragments among Kuwaiti individuals from the same cluster. Lower boxplots refer to TVD estimated on number of fragments, while upper boxplots refer to TVD estimated on length of fragments.

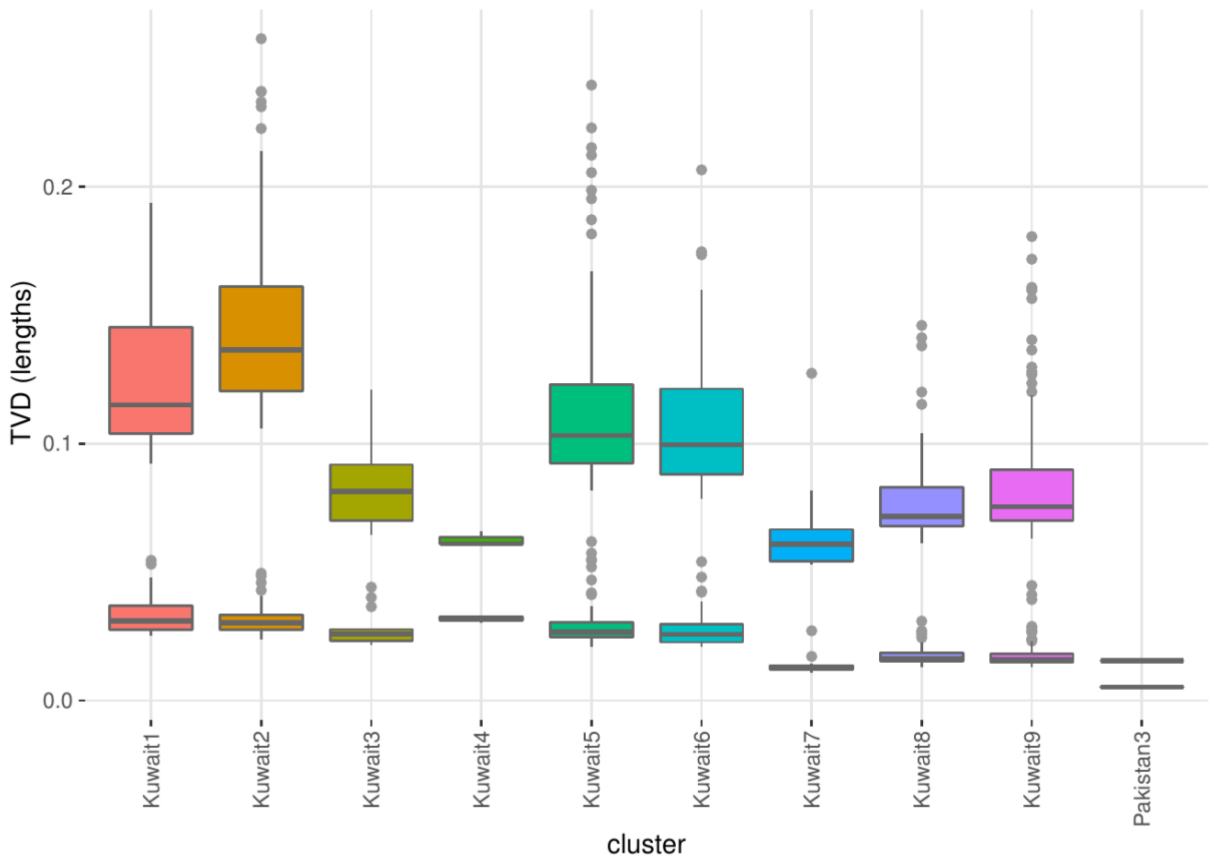

**Supplementary Fig. 7.** Ancestry proportions of the main eight Kuwait clusters as inferred by NNLS analysis using North/East Europe, Bedouins, Yoruba, Druze and North Africa clusters as putative sources.

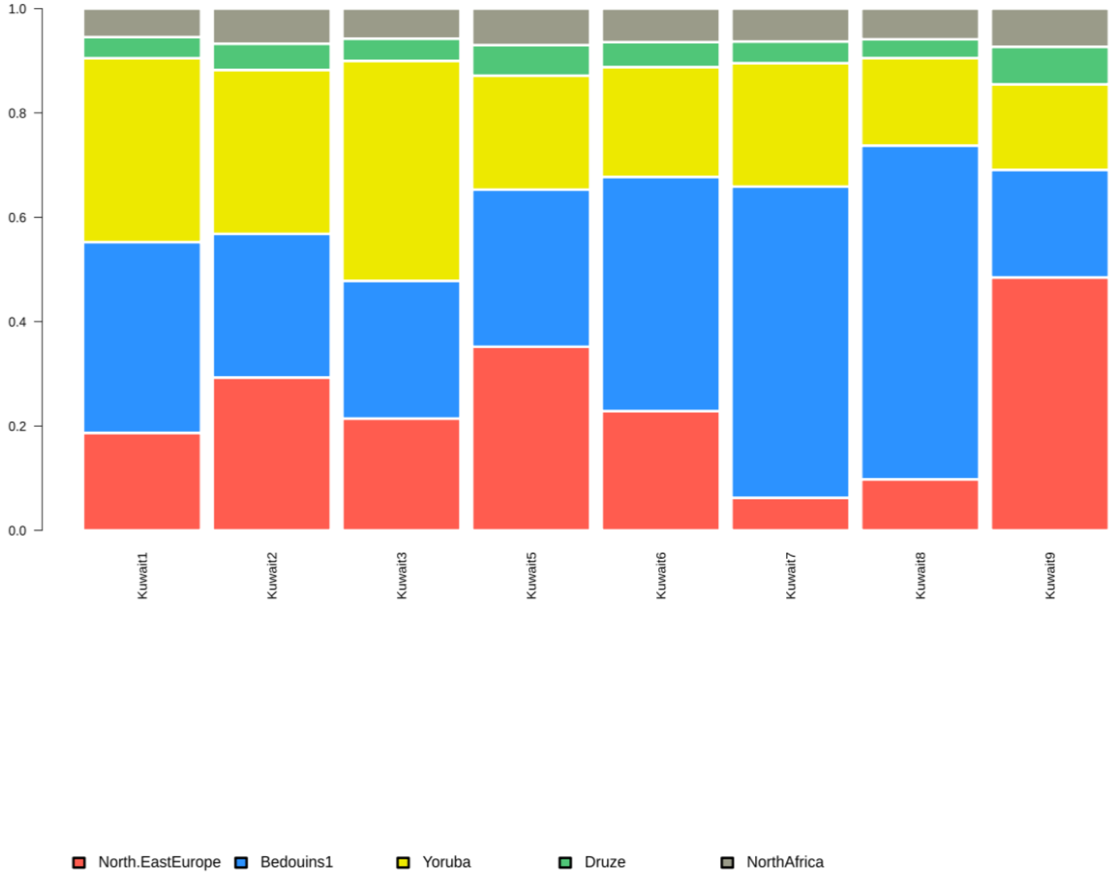

**Supplementary Fig. 8.** Inferred admixture sources by GLOBETROTTER population based analysis. (A) The x and y axes show the minor and major sources, respectively, as inferred by GLOBETROTTER. (B) Type of admixture.

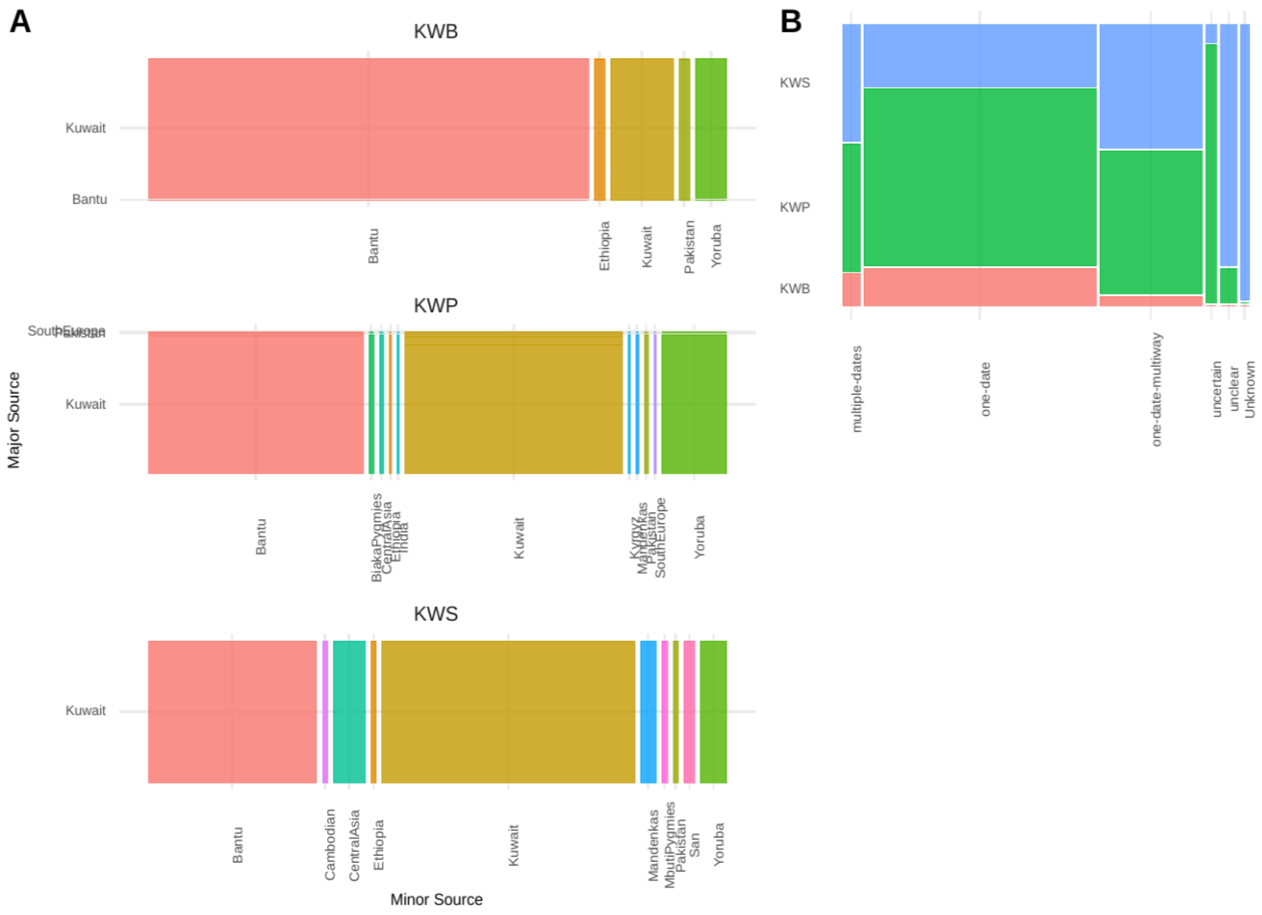

**Supplementary Table 1.** Details of the populations included in the study.

| Region | Population | No. of individuals | Publication |
|---|---|---|---|
| Middle East | Bedouin | 45 | Li et al 2008 |
| Middle East | Druze | 45 | Li et al 2008; Behar et al 2013 |
| Middle East | Lebanese | 8 | Behar et al 2010 |
| Middle East | Palestinian | 52 | Li et al 2008; Behar et al 2013 |
| Middle East | Samaritan | 2 | Behar et al 2010 |
| Middle East | Syrian | 16 | Behar et al 2010 |
| Middle East | Turkish | 19 | Behar et al 2010 |
| Middle East | Iranian | 19 | Behar et al 2010 |
| Middle East | Kurd | 6 | Yunusbayev et al 2012 |
| Middle East | Iranian Jewish | 12 | Behar et al 2010; Behar et al 2013 |
| Middle East | Iraqi Jewish | 13 | Behar et al 2010; Behar et al 2013 |
| Middle East | Kurdish Jewish | 9 | Behar et al. 2013 |
| Middle East | Syrian Jewish | 2 | Behar et al. 2013 |
| Middle East | Yemenite Jewish | 18 | Behar et al 2010; Behar et al 2013 |
| Arabian Peninsula | Kuwait | 583 | Present Study |
| Arabian Peninsula | Saudi | 20 | Behar et al 2010 |
| Arabian Peninsula | Yemeni | 8 | Behar et al 2010 |
| Arabian Peninsula | Jordanian | 20 | Behar et al 2010 |
| Sub-Saharan Africa | Bantu | 18 | Li et al 2008 |
| Central Africa | Biaka Pygmy | 22 | Li et al 2008 |
| North Africa | Egyptian | 12 | Behar et al 2010 |
| North Africa | Ethiopian | 19 | Behar et al 2010 |
| West Africa | Mandenka | 22 | Li et al 2008 |
| Central Africa | Mbuti Pygmy | 13 | Li et al 2008 |
| North Africa | Moroccan | 10 | Behar et al 2010 |
| North Africa | Mozabite | 27 | Li et al 2008 |
| South Africa | San | 5 | Li et al 2008 |
| West Africa | Yoruba | 21 | Li et al 2008 |
| South Caucasus | Abkhasian | 23 | Yunusbayev et al 2012; Behar et al 2013 |
| South Caucasus | Armenian | 16 | Yunusbayev et al 2012 |
| South Caucasus | Azeri | 16 | Yunusbayev et al 2015 |
| South Caucasus | Georgian | 30 | Behar et al 2010; Behar et al 2013 |
| North Caucasus | Adygei | 17 | Li et al 2008 |
| North Caucasus | Balkar | 22 | Yunusbayev et al 2015; Yunusbayev et al 2012 |
| North Caucasus | Chechen | 20 | Yunusbayev et al 2012 |
| North Caucasus | Kabardin | 3 | Yunusbayev et al 2015 |
| North Caucasus | Kumyk | 17 | Yunusbayev et al 2015; Yunusbayev et al 2012 |
| North Caucasus | Lezgin | 21 | Behar et al 2010; Behar et al 2013 |
| North Caucasus | Nogai | 16 | Yunusbayev et al 2012 |
| North Caucasus | North Ossetian | 18 | Yunusbayev et al 2012; Behar et al 2013 |
| North Caucasus | Tabasaran | 3 | Behar et al. 2013 |
| Central Asia | Kyrgyz | 20 | Yunusbayev et al 2015 |
| Central Asia | Tajik | 15 | Yunusbayev et al 2012 |
| Central Asia | Turkmen | 18 | Yunusbayev et al 2012; Yunusbayev et al 2015 |
| Central Asia | Uzbek | 19 | Behar et al 2010 |
| East Asia | Cambodian | 10 | Li et al 2008 |

| Region | Population | No. of individuals | Publication |
|---|---|---|---|
| East Asia | Han | 44 | Li et al 2008 |
| East Asia | Japanese | 28 | Li et al 2008 |
| East Asia | Uygur | 10 | Li et al 2008 |
| Western Europe | French | 28 | Li et al 2008 |
| Western Europe | French Basque | 24 | Li et al 2008 |
| Western Europe | Italian | 71 | Li et al 2008; Behar et al 2013 |
| Western Europe | Orcadian | 15 | Li et al 2008 |
| Western Europe | Spanish | 12 | Behar et al 2010 |
| Eastern Europe | Belarusian | 17 | Behar et al 2010; Behar et al 2013 |
| Eastern Europe | Bulgarian | 13 | Yunusbayev et al 2012 |
| Eastern Europe | Chuvash | 19 | Behar et al 2010; Yunusbayev et al 2015 |
| Eastern Europe | Croat | 24 | Behar et al. 2013 |
| Eastern Europe | Estonian | 15 | Raghavan et al. 2014 |
| Eastern Europe | Greek | 20 | Behar et al. 2013 |
| Eastern Europe | Hungarian | 19 | Behar et al 2010 |
| Eastern Europe | Lithuanian | 10 | Behar et al 2010 |
| Eastern Europe | Moldavian | 7 | Behar et al. 2013 |
| Eastern Europe | Mordovian | 15 | Yunusbayev et al 2012 |
| Eastern Europe | Polish | 17 | Behar et al. 2013 |
| Eastern Europe | Romanian | 16 | Behar et al 2010 |
| Eastern Europe | Russian | 23 | Yunusbayev et al 2015 |
| Eastern Europe | Swedish | 18 | Behar et al. 2013 |
| Eastern Europe | Tatar | 20 | Yunusbayev et al 2015 |
| Eastern Europe | Ukranian | 20 | Yunusbayev et al 2012 |
| Eastern Europe | Ashkenazi Jewish | 29 | Behar et al 2010; Behar et al 2013 |
| South Asia (Pakistan) | Balochi | 24 | Li et al 2008 |
| South Asia (Pakistan) | Brahui | 25 | Li et al 2008 |
| South Asia (Pakistan) | Burusho | 25 | Li et al 2008 |
| South Asia (Pakistan) | Hazara | 22 | Li et al 2008 |
| South Asia (Pakistan) | Kalash | 23 | Li et al 2008 |
| South Asia (Pakistan) | Makrani | 25 | Li et al 2008 |
| South Asia (Pakistan) | Pathan | 22 | Li et al 2008 |
| South Asia (Pakistan) | Sindhi | 24 | Li et al 2008 |
| South Asia (India) | Gujarati | 23 | Hapmap3 |
| South Asia (India) | Halakipikki | 4 | Metspalu et al 2011 |
| South Asia (India) | Malayan | 2 | Behar et al 2010 |
| South Asia (India) | North Kannadi | 8 | Behar et al 2010 |
| South Asia (India) | Paniya | 4 | Behar et al 2010 |
| South Asia (India) | Sakilli | 4 | Behar et al 2010 |

**Supplementary Table 2.** *$f_4$*-statistics comparing the genetic affinity of the Kuwait population subgroups to other modern populations.

| Pop1 | Pop2 | Pop3 | Pop4 | D val | std err | Z val |
|---|---|---|---|---|---|---|
| Kuwait-P | Kuwait-S | Gujaratis | Mbuti_Pygmies | 0.002097 | 0.000066 | 31.825 |
| Kuwait-P | Kuwait-S | Gujjar | Mbuti_Pygmies | 0.002043 | 0.000068 | 29.994 |
| Kuwait-P | Kuwait-S | Kshatriya | Mbuti_Pygmies | 0.002105 | 0.00007 | 29.985 |
| Kuwait-P | Kuwait-S | Pathan | Mbuti_Pygmies | 0.002001 | 0.000067 | 29.835 |
| Kuwait-P | Kuwait-S | Ror | Mbuti_Pygmies | 0.002052 | 0.000069 | 29.751 |
| Kuwait-P | Kuwait-S | Chamar | Mbuti_Pygmies | 0.002001 | 0.00007 | 28.733 |
| Kuwait-P | Kuwait-S | Gond | Mbuti_Pygmies | 0.001865 | 0.000067 | 27.647 |
| Kuwait-P | Kuwait-S | Pamiris | Mbuti_Pygmies | 0.001887 | 0.000068 | 27.564 |
| Kuwait-P | Kuwait-S | Brahmins_UP | Mbuti_Pygmies | 0.001982 | 0.000072 | 27.543 |
| Kuwait-P | Kuwait-S | Irula | Mbuti_Pygmies | 0.001898 | 0.00007 | 27.161 |
| Kuwait-P | Kuwait-S | Balochi | Mbuti_Pygmies | 0.001812 | 0.000067 | 27.065 |
| Kuwait-P | Kuwait-S | Paniya | Mbuti_Pygmies | 0.001864 | 0.000071 | 26.151 |
| Kuwait-P | Kuwait-S | Ho | Mbuti_Pygmies | 0.001778 | 0.000069 | 25.841 |
| Kuwait-P | Kuwait-S | Chukchis | Mbuti_Pygmies | 0.001838 | 0.000073 | 25.238 |
| Kuwait-P | Kuwait-S | Tajiks | Mbuti_Pygmies | 0.001747 | 0.000071 | 24.66 |
| Kuwait-P | Kuwait-S | Karitiana | Mbuti_Pygmies | 0.002152 | 0.000087 | 24.633 |
| Kuwait-P | Kuwait-S | Pashtun | Mbuti_Pygmies | 0.001816 | 0.000074 | 24.599 |
| Kuwait-P | Kuwait-S | Kusundas | Mbuti_Pygmies | 0.001787 | 0.000073 | 24.317 |
| Kuwait-P | Kuwait-S | Kyrgyzians | Mbuti_Pygmies | 0.001679 | 0.000071 | 23.675 |
| Kuwait-P | Kuwait-S | Uygurs | Mbuti_Pygmies | 0.001716 | 0.000073 | 23.505 |
| Kuwait-P | Kuwait-S | Lezgins | Mbuti_Pygmies | 0.001566 | 0.00007 | 22.221 |
| Kuwait-P | Kuwait-S | Japanese | Mbuti_Pygmies | 0.001652 | 0.000075 | 21.894 |
| Kuwait-P | Kuwait-S | Han | Mbuti_Pygmies | 0.001646 | 0.000075 | 21.887 |
| Kuwait-P | Kuwait-S | Chuvash | Mbuti_Pygmies | 0.001561 | 0.000072 | 21.551 |
| Kuwait-P | Kuwait-S | Cambodians | Mbuti_Pygmies | 0.001599 | 0.000075 | 21.312 |
| Kuwait-P | Kuwait-S | Turkmens | Mbuti_Pygmies | 0.001458 | 0.000069 | 21.236 |
| Kuwait-P | Kuwait-S | Tatars | Mbuti_Pygmies | 0.001496 | 0.000072 | 20.877 |
| Kuwait-P | Kuwait-S | Chechens | Mbuti_Pygmies | 0.001478 | 0.000071 | 20.7 |
| Kuwait-P | Kuwait-S | Nogais | Mbuti_Pygmies | 0.001434 | 0.000072 | 20.049 |
| Kuwait-P | Kuwait-S | Kumyks | Mbuti_Pygmies | 0.001391 | 0.00007 | 19.972 |
| Kuwait-P | Kuwait-S | Parsis | Mbuti_Pygmies | 0.001394 | 0.00007 | 19.863 |
| Kuwait-P | Kuwait-S | Balkars | Mbuti_Pygmies | 0.001398 | 0.00007 | 19.848 |
| Kuwait-P | Kuwait-S | Onge | Mbuti_Pygmies | 0.001538 | 0.000081 | 19.083 |
| Kuwait-P | Kuwait-S | Tabasaran | Mbuti_Pygmies | 0.001576 | 0.000084 | 18.711 |
| Kuwait-P | Kuwait-S | Abkhasians | Mbuti_Pygmies | 0.001314 | 0.00007 | 18.684 |
| Kuwait-P | Kuwait-S | Kabardins | Mbuti_Pygmies | 0.00145 | 0.00008 | 18.123 |
| Kuwait-P | Kuwait-S | Ukranians | Mbuti_Pygmies | 0.001271 | 0.000073 | 17.396 |
| Kuwait-P | Kuwait-S | Polish | Mbuti_Pygmies | 0.001297 | 0.000075 | 17.265 |
| Kuwait-P | Kuwait-S | Russians | Mbuti_Pygmies | 0.001347 | 0.000078 | 17.201 |
| Kuwait-P | Kuwait-S | Georgians | Mbuti_Pygmies | 0.001216 | 0.000073 | 16.596 |

| Pop1 | Pop2 | Pop3 | Pop4 | D val | std err | Z val |
|---|---|---|---|---|---|---|
| Kuwait-P | Kuwait-S | Moldavians | Mbuti_Pygmies | 0.001275 | 0.000077 | 16.582 |
| Kuwait-P | Kuwait-S | Azeris | Mbuti_Pygmies | 0.001153 | 0.00007 | 16.557 |
| Kuwait-P | Kuwait-S | Orcadians | Mbuti_Pygmies | 0.001231 | 0.000074 | 16.556 |
| Kuwait-P | Kuwait-S | Hungarians | Mbuti_Pygmies | 0.001148 | 0.000074 | 15.494 |
| Kuwait-P | Kuwait-S | Turks | Mbuti_Pygmies | 0.001076 | 0.00007 | 15.449 |
| Kuwait-P | Kuwait-S | Kurds | Mbuti_Pygmies | 0.001146 | 0.000075 | 15.254 |
| Kuwait-P | Kuwait-S | Iranians | Mbuti_Pygmies | 0.001045 | 0.000069 | 15.202 |
| Kuwait-P | Kuwait-S | Croaitians | Mbuti_Pygmies | 0.001088 | 0.000073 | 14.828 |
| Kuwait-P | Kuwait-S | Romanians | Mbuti_Pygmies | 0.00105 | 0.000073 | 14.387 |
| Kuwait-P | Kuwait-S | Romanians | Mbuti_Pygmies | 0.00105 | 0.000073 | 14.387 |
| Kuwait-P | Kuwait-S | Armenians | Mbuti_Pygmies | 0.000974 | 0.00007 | 14.008 |
| Kuwait-P | Kuwait-S | Greeks | Mbuti_Pygmies | 0.000848 | 0.000073 | 11.621 |
| Kuwait-P | Kuwait-S | FrenchBasques | Mbuti_Pygmies | 0.000831 | 0.000074 | 11.187 |
| Kuwait-P | Kuwait-S | Spaniards | Mbuti_Pygmies | 0.000832 | 0.000075 | 11.083 |
| Kuwait-P | Kuwait-S | Tuscans | Mbuti_Pygmies | 0.00081 | 0.000075 | 10.808 |
| Kuwait-P | Kuwait-S | North_Italians | Mbuti_Pygmies | 0.000803 | 0.000077 | 10.458 |
| Kuwait-P | Kuwait-S | KurdishJews | Mbuti_Pygmies | 0.000714 | 0.000072 | 9.893 |
| Kuwait-P | Kuwait-S | IranianJewish | Mbuti_Pygmies | 0.000674 | 0.00007 | 9.66 |
| Kuwait-P | Kuwait-S | AshkenaziJewish | Mbuti_Pygmies | 0.000625 | 0.00007 | 8.92 |
| Kuwait-P | Kuwait-S | IraqiJewish | Mbuti_Pygmies | 0.00058 | 0.000071 | 8.228 |
| Kuwait-P | Kuwait-S | Druze | Mbuti_Pygmies | 0.00044 | 0.000068 | 6.467 |
| Kuwait-P | Kuwait-S | SyrianJews | Mbuti_Pygmies | 0.000526 | 0.000091 | 5.752 |
| Kuwait-P | Kuwait-S | Sardinians | Mbuti_Pygmies | 0.000402 | 0.000073 | 5.5 |
| Kuwait-P | Kuwait-S | Syrians | Mbuti_Pygmies | 0.000331 | 0.00007 | 4.715 |
| Kuwait-P | Kuwait-S | Lebanese | Mbuti_Pygmies | 0.000302 | 0.000072 | 4.185 |
| Kuwait-P | Kuwait-S | Jordanians | Mbuti_Pygmies | 0.000093 | 0.000064 | 1.463 |
| Kuwait-P | Kuwait-S | Palestinians | Mbuti_Pygmies | 0.000054 | 0.000064 | 0.843 |
| Kuwait-P | Kuwait-S | Samartians | Mbuti_Pygmies | 0.000064 | 0.000091 | 0.708 |
| **Kuwait-P** | **Kuwait-S** | **Mandenka** | **Mbuti_Pygmies** | **-0.00006** | **0.000035** | **-1.716** |
| **Kuwait-P** | **Kuwait-S** | **Mozabites** | **Mbuti_Pygmies** | **-0.000295** | **0.000061** | **-4.825** |
| **Kuwait-P** | **Kuwait-S** | **Egyptians** | **Mbuti_Pygmies** | **-0.000351** | **0.000064** | **-5.505** |
| **Kuwait-P** | **Kuwait-S** | **Yemenese** | **Mbuti_Pygmies** | **-0.000448** | **0.000063** | **-7.12** |
| **Kuwait-P** | **Kuwait-S** | **Moroccans** | **Mbuti_Pygmies** | **-0.000436** | **0.000061** | **-7.131** |
| **Kuwait-P** | **Kuwait-S** | **Kuwait-B** | **Mbuti_Pygmies** | **-0.000412** | **0.000051** | **-8.135** |
| **Kuwait-P** | **Kuwait-S** | **YemeniteJews** | **Mbuti_Pygmies** | **-0.000767** | **0.000068** | **-11.259** |
| **Kuwait-P** | **Kuwait-S** | **Bedouins** | **Mbuti_Pygmies** | **-0.000936** | **0.000065** | **-14.426** |
| **Kuwait-P** | **Kuwait-S** | **Ethiopians** | **Mbuti_Pygmies** | **-0.000865** | **0.000047** | **-18.392** |
| **Kuwait-P** | **Kuwait-S** | **Saudis** | **Mbuti_Pygmies** | **-0.001409** | **0.000067** | **-21.098** |
| Kuwait-B | Kuwait-S | Mandenka | Mbuti_Pygmies | -0.000501 | 0.000055 | -9.063 |
| Kuwait-B | Kuwait-S | Onge | Mbuti_Pygmies | -0.008888 | 0.000134 | -66.54 |
| Kuwait-B | Kuwait-S | Karitiana | Mbuti_Pygmies | -0.009863 | 0.000145 | -67.986 |
| Kuwait-B | Kuwait-S | Japanese | Mbuti_Pygmies | -0.00932 | 0.000136 | -68.456 |
| Kuwait-B | Kuwait-S | Cambodians | Mbuti_Pygmies | -0.009241 | 0.000132 | -70.22 |

| Pop1 | Pop2 | Pop3 | Pop4 | D val | std err | Z val |
|---|---|---|---|---|---|---|
| Kuwait-B | Kuwait-S | Han | Mbuti_Pygmies | -0.009316 | 0.000132 | -70.68 |
| Kuwait-B | Kuwait-S | Kusundas | Mbuti_Pygmies | -0.009414 | 0.000132 | -71.079 |
| Kuwait-B | Kuwait-S | Paniya | Mbuti_Pygmies | -0.009172 | 0.000127 | -72.037 |
| Kuwait-B | Kuwait-S | Ethiopians | Mbuti_Pygmies | -0.006282 | 0.000087 | -72.054 |
| Kuwait-B | Kuwait-S | Irula | Mbuti_Pygmies | -0.00942 | 0.000128 | -73.339 |
| Kuwait-B | Kuwait-S | Ho | Mbuti_Pygmies | -0.00919 | 0.000125 | -73.757 |
| Kuwait-B | Kuwait-S | Gond | Mbuti_Pygmies | -0.009306 | 0.000125 | -74.349 |
| Kuwait-B | Kuwait-S | Chamar | Mbuti_Pygmies | -0.009652 | 0.000129 | -74.883 |
| Kuwait-B | Kuwait-S | Chukchis | Mbuti_Pygmies | -0.009999 | 0.000133 | -75.133 |
| Kuwait-B | Kuwait-S | Kyrgyzians | Mbuti_Pygmies | -0.010047 | 0.000132 | -76.383 |
| Kuwait-B | Kuwait-S | Uygurs | Mbuti_Pygmies | -0.010231 | 0.000133 | -76.971 |
| Kuwait-B | Kuwait-S | Kshatriya | Mbuti_Pygmies | -0.010103 | 0.000131 | -76.977 |
| Kuwait-B | Kuwait-S | SyrianJews | Mbuti_Pygmies | -0.011594 | 0.00015 | -77.081 |
| Kuwait-B | Kuwait-S | Samartians | Mbuti_Pygmies | -0.011887 | 0.000152 | -78.248 |
| Kuwait-B | Kuwait-S | Moroccans | Mbuti_Pygmies | -0.008719 | 0.000111 | -78.878 |
| Kuwait-B | Kuwait-S | Brahmins_UP | Mbuti_Pygmies | -0.010208 | 0.000129 | -79.085 |
| Kuwait-B | Kuwait-S | Gujjar | Mbuti_Pygmies | -0.010316 | 0.00013 | -79.283 |
| Kuwait-B | Kuwait-S | Tabasaran | Mbuti_Pygmies | -0.011385 | 0.000143 | -79.394 |
| Kuwait-B | Kuwait-S | Yemenese | Mbuti_Pygmies | -0.00966 | 0.000121 | -79.567 |
| Kuwait-B | Kuwait-S | Kabardins | Mbuti_Pygmies | -0.011369 | 0.000142 | -80.001 |
| Kuwait-B | Kuwait-S | Gujaratis | Mbuti_Pygmies | -0.010037 | 0.000125 | -80.145 |
| Kuwait-B | Kuwait-S | Pashtun | Mbuti_Pygmies | -0.010743 | 0.000133 | -80.662 |
| Kuwait-B | Kuwait-S | Mozabites | Mbuti_Pygmies | -0.009262 | 0.000113 | -81.633 |
| Kuwait-B | Kuwait-S | Ror | Mbuti_Pygmies | -0.010548 | 0.000128 | -82.26 |
| Kuwait-B | Kuwait-S | Balochi | Mbuti_Pygmies | -0.010457 | 0.000127 | -82.485 |
| Kuwait-B | Kuwait-S | Chuvash | Mbuti_Pygmies | -0.011017 | 0.000133 | -82.681 |
| Kuwait-B | Kuwait-S | Tajiks | Mbuti_Pygmies | -0.010834 | 0.000131 | -82.716 |
| Kuwait-B | Kuwait-S | Nogais | Mbuti_Pygmies | -0.011097 | 0.000134 | -82.963 |
| Kuwait-B | Kuwait-S | Moldavians | Mbuti_Pygmies | -0.011531 | 0.000139 | -83.083 |
| Kuwait-B | Kuwait-S | Pathan | Mbuti_Pygmies | -0.010551 | 0.000127 | -83.32 |
| Kuwait-B | Kuwait-S | Parsis | Mbuti_Pygmies | -0.011009 | 0.000132 | -83.586 |
| Kuwait-B | Kuwait-S | Turkmens | Mbuti_Pygmies | -0.010855 | 0.00013 | -83.641 |
| Kuwait-B | Kuwait-S | Kurds | Mbuti_Pygmies | -0.011526 | 0.000137 | -84.08 |
| Kuwait-B | Kuwait-S | Tatars | Mbuti_Pygmies | -0.01113 | 0.000132 | -84.224 |
| Kuwait-B | Kuwait-S | Pamiris | Mbuti_Pygmies | -0.010906 | 0.000129 | -84.264 |
| Kuwait-B | Kuwait-S | Iranians | Mbuti_Pygmies | -0.011083 | 0.000131 | -84.353 |
| Kuwait-B | Kuwait-S | Polish | Mbuti_Pygmies | -0.011638 | 0.000138 | -84.475 |
| Kuwait-B | Kuwait-S | Tuscans | Mbuti_Pygmies | -0.011816 | 0.00014 | -84.622 |
| Kuwait-B | Kuwait-S | Russians | Mbuti_Pygmies | -0.011592 | 0.000137 | -84.646 |
| Kuwait-B | Kuwait-S | Egyptians | Mbuti_Pygmies | -0.009995 | 0.000118 | -84.763 |
| Kuwait-B | Kuwait-S | Balkars | Mbuti_Pygmies | -0.01136 | 0.000134 | -84.843 |
| Kuwait-B | Kuwait-S | Lezgins | Mbuti_Pygmies | -0.01141 | 0.000134 | -85.027 |
| Kuwait-B | Kuwait-S | KurdishJews | Mbuti_Pygmies | -0.011665 | 0.000137 | -85.342 |

| Pop1 | Pop2 | Pop3 | Pop4 | D val | std err | Z val |
|---|---|---|---|---|---|---|
| Kuwait-B | Kuwait-S | IranianJewish | Mbuti_Pygmies | -0.011606 | 0.000136 | -85.412 |
| Kuwait-B | Kuwait-S | Hungarians | Mbuti_Pygmies | -0.011657 | 0.000136 | -85.432 |
| Kuwait-B | Kuwait-S | Orcadians | Mbuti_Pygmies | -0.011627 | 0.000136 | -85.464 |
| Kuwait-B | Kuwait-S | Georgians | Mbuti_Pygmies | -0.011563 | 0.000135 | -85.587 |
| Kuwait-B | Kuwait-S | Chechens | Mbuti_Pygmies | -0.011447 | 0.000134 | -85.632 |
| Kuwait-B | Kuwait-S | Spaniards | Mbuti_Pygmies | -0.011667 | 0.000135 | -86.285 |
| Kuwait-B | Kuwait-S | FrenchBasques | Mbuti_Pygmies | -0.011834 | 0.000137 | -86.288 |
| Kuwait-B | Kuwait-S | Ukranians | Mbuti_Pygmies | -0.011537 | 0.000134 | -86.288 |
| Kuwait-B | Kuwait-S | Kumyks | Mbuti_Pygmies | -0.011396 | 0.000132 | -86.34 |
| Kuwait-B | Kuwait-S | Turks | Mbuti_Pygmies | -0.011456 | 0.000133 | -86.435 |
| Kuwait-B | Kuwait-S | Croaitians | Mbuti_Pygmies | -0.011706 | 0.000135 | -86.464 |
| Kuwait-B | Kuwait-S | Abkhasians | Mbuti_Pygmies | -0.011587 | 0.000134 | -86.698 |
| Kuwait-B | Kuwait-S | Jordanians | Mbuti_Pygmies | -0.010831 | 0.000125 | -86.718 |
| Kuwait-B | Kuwait-S | Lebanese | Mbuti_Pygmies | -0.011318 | 0.00013 | -86.741 |
| Kuwait-B | Kuwait-S | Azeris | Mbuti_Pygmies | -0.011366 | 0.000131 | -86.891 |
| Kuwait-B | Kuwait-S | Greeks | Mbuti_Pygmies | -0.011785 | 0.000135 | -87.056 |
| Kuwait-B | Kuwait-S | Romanians | Mbuti_Pygmies | -0.011655 | 0.000134 | -87.08 |
| Kuwait-B | Kuwait-S | Romanians | Mbuti_Pygmies | -0.011655 | 0.000134 | -87.08 |
| Kuwait-B | Kuwait-S | North_Italians | Mbuti_Pygmies | -0.011843 | 0.000136 | -87.374 |
| Kuwait-B | Kuwait-S | Syrians | Mbuti_Pygmies | -0.011289 | 0.000128 | -88.09 |
| Kuwait-B | Kuwait-S | IraqiJewish | Mbuti_Pygmies | -0.011731 | 0.000133 | -88.443 |
| Kuwait-B | Kuwait-S | Armenians | Mbuti_Pygmies | -0.011674 | 0.000132 | -88.492 |
| Kuwait-B | Kuwait-S | Sardinians | Mbuti_Pygmies | -0.011996 | 0.000135 | -88.766 |
| Kuwait-B | Kuwait-S | Palestinians | Mbuti_Pygmies | -0.011064 | 0.000125 | -88.795 |
| Kuwait-B | Kuwait-S | AshkenaziJewish | Mbuti_Pygmies | -0.011639 | 0.00013 | -89.375 |
| Kuwait-B | Kuwait-S | Druze | Mbuti_Pygmies | -0.011646 | 0.00013 | -89.611 |
| Kuwait-B | Kuwait-S | YemeniteJews | Mbuti_Pygmies | -0.011786 | 0.000128 | -92.173 |
| Kuwait-B | Kuwait-S | Bedouins | Mbuti_Pygmies | -0.011496 | 0.000123 | -93.306 |
| Kuwait-B | Kuwait-S | Saudis | Mbuti_Pygmies | -0.012053 | 0.000128 | -93.91 |
| Kuwait-B | Kuwait-P | Mandenka | Mbuti_Pygmies | -0.000441 | 0.000055 | -8.064 |
| Kuwait-B | Kuwait-P | Ethiopians | Mbuti_Pygmies | -0.005417 | 0.000085 | -63.614 |
| Kuwait-B | Kuwait-P | Moroccans | Mbuti_Pygmies | -0.008283 | 0.000113 | -73.46 |
| Kuwait-B | Kuwait-P | Onge | Mbuti_Pygmies | -0.010426 | 0.000142 | -73.638 |
| Kuwait-B | Kuwait-P | Yemenese | Mbuti_Pygmies | -0.009212 | 0.000123 | -74.944 |
| Kuwait-B | Kuwait-P | Samartians | Mbuti_Pygmies | -0.011951 | 0.000156 | -76.491 |
| Kuwait-B | Kuwait-P | Karitiana | Mbuti_Pygmies | -0.012015 | 0.000157 | -76.655 |
| Kuwait-B | Kuwait-P | Japanese | Mbuti_Pygmies | -0.010972 | 0.000142 | -77.11 |
| Kuwait-B | Kuwait-P | Mozabites | Mbuti_Pygmies | -0.008967 | 0.000116 | -77.333 |
| Kuwait-B | Kuwait-P | Cambodians | Mbuti_Pygmies | -0.01084 | 0.000139 | -78.224 |
| Kuwait-B | Kuwait-P | Han | Mbuti_Pygmies | -0.010962 | 0.00014 | -78.328 |
| Kuwait-B | Kuwait-P | SyrianJews | Mbuti_Pygmies | -0.01212 | 0.000154 | -78.912 |
| Kuwait-B | Kuwait-P | Egyptians | Mbuti_Pygmies | -0.009644 | 0.000119 | -80.845 |
| Kuwait-B | Kuwait-P | Kusundas | Mbuti_Pygmies | -0.011201 | 0.000139 | -80.861 |

| Pop1 | Pop2 | Pop3 | Pop4 | D val | std err | Z val |
|---|---|---|---|---|---|---|
| Kuwait-B | Kuwait-P | Paniya | Mbuti_Pygmies | -0.011036 | 0.000135 | -81.863 |
| Kuwait-B | Kuwait-P | Saudis | Mbuti_Pygmies | -0.010644 | 0.00013 | -82.119 |
| Kuwait-B | Kuwait-P | Ho | Mbuti_Pygmies | -0.010967 | 0.000133 | -82.425 |
| Kuwait-B | Kuwait-P | Kyrgyzians | Mbuti_Pygmies | -0.011725 | 0.000139 | -84.084 |
| Kuwait-B | Kuwait-P | Chukchis | Mbuti_Pygmies | -0.011837 | 0.000141 | -84.129 |
| Kuwait-B | Kuwait-P | Irula | Mbuti_Pygmies | -0.011318 | 0.000134 | -84.25 |
| Kuwait-B | Kuwait-P | Gond | Mbuti_Pygmies | -0.011172 | 0.000132 | -84.351 |
| Kuwait-B | Kuwait-P | Bedouins | Mbuti_Pygmies | -0.01056 | 0.000125 | -84.816 |
| Kuwait-B | Kuwait-P | YemeniteJews | Mbuti_Pygmies | -0.011019 | 0.000129 | -85.607 |
| Kuwait-B | Kuwait-P | Chamar | Mbuti_Pygmies | -0.011653 | 0.000136 | -85.705 |
| Kuwait-B | Kuwait-P | Jordanians | Mbuti_Pygmies | -0.010924 | 0.000127 | -85.853 |
| Kuwait-B | Kuwait-P | Uygurs | Mbuti_Pygmies | -0.011947 | 0.000139 | -86.058 |
| Kuwait-B | Kuwait-P | Kabardins | Mbuti_Pygmies | -0.012819 | 0.000149 | -86.089 |
| Kuwait-B | Kuwait-P | Palestinians | Mbuti_Pygmies | -0.011118 | 0.000128 | -86.667 |
| Kuwait-B | Kuwait-P | Tuscans | Mbuti_Pygmies | -0.012625 | 0.000145 | -87.229 |
| Kuwait-B | Kuwait-P | Lebanese | Mbuti_Pygmies | -0.01162 | 0.000133 | -87.458 |
| Kuwait-B | Kuwait-P | Tabasaran | Mbuti_Pygmies | -0.012961 | 0.000148 | -87.782 |
| Kuwait-B | Kuwait-P | IranianJewish | Mbuti_Pygmies | -0.01228 | 0.00014 | -87.816 |
| Kuwait-B | Kuwait-P | Brahmins_UP | Mbuti_Pygmies | -0.01219 | 0.000139 | -88.003 |
| Kuwait-B | Kuwait-P | Kshatriya | Mbuti_Pygmies | -0.012208 | 0.000139 | -88.097 |
| Kuwait-B | Kuwait-P | Kurds | Mbuti_Pygmies | -0.012672 | 0.000144 | -88.109 |
| Kuwait-B | Kuwait-P | Pashtun | Mbuti_Pygmies | -0.012559 | 0.000142 | -88.154 |
| Kuwait-B | Kuwait-P | KurdishJews | Mbuti_Pygmies | -0.012379 | 0.00014 | -88.242 |
| Kuwait-B | Kuwait-P | Syrians | Mbuti_Pygmies | -0.01162 | 0.000132 | -88.28 |
| Kuwait-B | Kuwait-P | Moldavians | Mbuti_Pygmies | -0.012807 | 0.000145 | -88.384 |
| Kuwait-B | Kuwait-P | Sardinians | Mbuti_Pygmies | -0.012397 | 0.00014 | -88.645 |
| Kuwait-B | Kuwait-P | Nogais | Mbuti_Pygmies | -0.012531 | 0.000141 | -88.856 |
| Kuwait-B | Kuwait-P | Spaniards | Mbuti_Pygmies | -0.012499 | 0.00014 | -89.173 |
| Kuwait-B | Kuwait-P | FrenchBasques | Mbuti_Pygmies | -0.012665 | 0.000142 | -89.185 |
| Kuwait-B | Kuwait-P | Polish | Mbuti_Pygmies | -0.012935 | 0.000145 | -89.228 |
| Kuwait-B | Kuwait-P | IraqiJewish | Mbuti_Pygmies | -0.012311 | 0.000137 | -89.533 |
| Kuwait-B | Kuwait-P | Iranians | Mbuti_Pygmies | -0.012128 | 0.000135 | -89.571 |
| Kuwait-B | Kuwait-P | Orcadians | Mbuti_Pygmies | -0.012858 | 0.000143 | -89.712 |
| Kuwait-B | Kuwait-P | Druze | Mbuti_Pygmies | -0.012086 | 0.000135 | -89.723 |
| Kuwait-B | Kuwait-P | Greeks | Mbuti_Pygmies | -0.012634 | 0.000141 | -89.832 |
| Kuwait-B | Kuwait-P | Gujjar | Mbuti_Pygmies | -0.012359 | 0.000138 | -89.881 |
| Kuwait-B | Kuwait-P | Chuvash | Mbuti_Pygmies | -0.012578 | 0.00014 | -89.936 |
| Kuwait-B | Kuwait-P | Parsis | Mbuti_Pygmies | -0.012404 | 0.000138 | -90.113 |
| Kuwait-B | Kuwait-P | Russians | Mbuti_Pygmies | -0.01294 | 0.000143 | -90.285 |
| Kuwait-B | Kuwait-P | Romanians | Mbuti_Pygmies | -0.012705 | 0.00014 | -90.436 |
| Kuwait-B | Kuwait-P | Romanians | Mbuti_Pygmies | -0.012705 | 0.00014 | -90.436 |
| Kuwait-B | Kuwait-P | Hungarians | Mbuti_Pygmies | -0.012805 | 0.000141 | -90.503 |
| Kuwait-B | Kuwait-P | Turkmens | Mbuti_Pygmies | -0.012312 | 0.000136 | -90.514 |

| Pop1 | Pop2 | Pop3 | Pop4 | D val | std err | Z val |
|---|---|---|---|---|---|---|
| Kuwait-B | Kuwait-P | North_Italians | Mbuti_Pygmies | -0.012647 | 0.000139 | -90.69 |
| Kuwait-B | Kuwait-P | Tajiks | Mbuti_Pygmies | -0.012581 | 0.000138 | -90.864 |
| Kuwait-B | Kuwait-P | Tatars | Mbuti_Pygmies | -0.012625 | 0.000139 | -90.888 |
| Kuwait-B | Kuwait-P | Croaitians | Mbuti_Pygmies | -0.012793 | 0.000141 | -90.915 |
| Kuwait-B | Kuwait-P | AshkenaziJewish | Mbuti_Pygmies | -0.012264 | 0.000135 | -90.989 |
| Kuwait-B | Kuwait-P | Azeris | Mbuti_Pygmies | -0.012519 | 0.000137 | -91.123 |
| Kuwait-B | Kuwait-P | Turks | Mbuti_Pygmies | -0.012532 | 0.000137 | -91.164 |
| Kuwait-B | Kuwait-P | Georgians | Mbuti_Pygmies | -0.012779 | 0.00014 | -91.253 |
| Kuwait-B | Kuwait-P | Ukranians | Mbuti_Pygmies | -0.012808 | 0.00014 | -91.38 |
| Kuwait-B | Kuwait-P | Gujaratis | Mbuti_Pygmies | -0.012134 | 0.000133 | -91.409 |
| Kuwait-B | Kuwait-P | Chechens | Mbuti_Pygmies | -0.012925 | 0.000141 | -91.722 |
| Kuwait-B | Kuwait-P | Abkhasians | Mbuti_Pygmies | -0.012901 | 0.000141 | -91.771 |
| Kuwait-B | Kuwait-P | Balkars | Mbuti_Pygmies | -0.012757 | 0.000139 | -91.999 |
| Kuwait-B | Kuwait-P | Balochi | Mbuti_Pygmies | -0.012268 | 0.000133 | -92.002 |
| Kuwait-B | Kuwait-P | Kumyks | Mbuti_Pygmies | -0.012787 | 0.000138 | -92.381 |
| Kuwait-B | Kuwait-P | Ror | Mbuti_Pygmies | -0.0126 | 0.000136 | -92.446 |
| Kuwait-B | Kuwait-P | Lezgins | Mbuti_Pygmies | -0.012976 | 0.00014 | -92.468 |
| Kuwait-B | Kuwait-P | Armenians | Mbuti_Pygmies | -0.012648 | 0.000137 | -92.55 |
| Kuwait-B | Kuwait-P | Pathan | Mbuti_Pygmies | -0.012552 | 0.000135 | -92.939 |
| Kuwait-B | Kuwait-P | Pamiris | Mbuti_Pygmies | -0.012793 | 0.000138 | -93.036 |

**Supplementary Table 3.** *$f_4$*-statistics for relative allele sharing of the Kuwait population subgroups to ancient West Eurasian individuals.

| Pop1 | Pop2 | Pop3 | Pop4 | D val | std err | Z val |
|---|---|---|---|---|---|---|
| Kuwait-B | Kuwait-P | Iran_HotuIIIb | Mbuti_Pygmies | -0.012446 | 0.000298 | -41.718 |
| Kuwait-B | Kuwait-P | Natufian | Mbuti_Pygmies | -0.009737 | 0.000185 | -52.651 |
| Kuwait-B | Kuwait-P | Ust_Ishim | Mbuti_Pygmies | -0.009782 | 0.000178 | -54.863 |
| Kuwait-B | Kuwait-P | Iberia_BA | Mbuti_Pygmies | -0.012631 | 0.000226 | -55.838 |
| Kuwait-B | Kuwait-P | Kostenki14 | Mbuti_Pygmies | -0.011138 | 0.000183 | -60.786 |
| Kuwait-B | Kuwait-P | Levant_N | Mbuti_Pygmies | -0.011009 | 0.00017 | -64.813 |
| Kuwait-B | Kuwait-P | Anatolia_ChL | Mbuti_Pygmies | -0.012736 | 0.000196 | -64.867 |
| Kuwait-B | Kuwait-P | MA1 | Mbuti_Pygmies | -0.012685 | 0.000188 | -67.412 |
| Kuwait-B | Kuwait-P | Clovis_Anzick | Mbuti_Pygmies | -0.011746 | 0.000174 | -67.692 |
| Kuwait-B | Kuwait-P | Switzerland_HG | Mbuti_Pygmies | -0.01231 | 0.00018 | -68.305 |
| Kuwait-B | Kuwait-P | Anatolia_Ottoman | Mbuti_Pygmies | -0.011998 | 0.000175 | -68.744 |
| Kuwait-B | Kuwait-P | Anatolia_IA | Mbuti_Pygmies | -0.012334 | 0.000178 | -69.484 |
| Kuwait-B | Kuwait-P | Iran_recent | Mbuti_Pygmies | -0.012641 | 0.000181 | -69.705 |
| Kuwait-B | Kuwait-P | Steppe_IA | Mbuti_Pygmies | -0.012901 | 0.000182 | -70.812 |
| Kuwait-B | Kuwait-P | Yamnaya_EBA | Mbuti_Pygmies | -0.013176 | 0.000185 | -71.137 |
| Kuwait-B | Kuwait-P | Turkmenistan_IA | Mbuti_Pygmies | -0.012874 | 0.000176 | -73.073 |
| Kuwait-B | Kuwait-P | Levant_BA | Mbuti_Pygmies | -0.011464 | 0.000156 | -73.327 |
| Kuwait-B | Kuwait-P | Iran_N | Mbuti_Pygmies | -0.012524 | 0.000171 | -73.413 |
| Kuwait-B | Kuwait-P | WestSiberia_HG | Mbuti_Pygmies | -0.013052 | 0.000174 | -74.803 |
| Kuwait-B | Kuwait-P | Indus_Diaspora | Mbuti_Pygmies | -0.012226 | 0.000159 | -76.757 |
| Kuwait-B | Kuwait-P | EHG | Mbuti_Pygmies | -0.013078 | 0.000169 | -77.256 |
| Kuwait-B | Kuwait-P | WHG | Mbuti_Pygmies | -0.012308 | 0.000159 | -77.528 |
| Kuwait-B | Kuwait-P | Namazga_CA | Mbuti_Pygmies | -0.012792 | 0.000162 | -79.196 |
| Kuwait-B | Kuwait-P | CHG | Mbuti_Pygmies | -0.012969 | 0.000161 | -80.477 |
| Kuwait-B | Kuwait-P | SHG | Mbuti_Pygmies | -0.012701 | 0.000157 | -80.681 |
| Kuwait-B | Kuwait-P | Anatolia_EBA | Mbuti_Pygmies | -0.012219 | 0.00015 | -81.311 |
| Kuwait-B | Kuwait-P | Armenia_EBA | Mbuti_Pygmies | -0.012849 | 0.000157 | -82.024 |
| Kuwait-B | Kuwait-P | Steppe_LBA | Mbuti_Pygmies | -0.012714 | 0.000154 | -82.311 |
| Kuwait-B | Kuwait-P | Armenia_ChL | Mbuti_Pygmies | -0.012733 | 0.000154 | -82.448 |
| Kuwait-B | Kuwait-P | IranTuran_N | Mbuti_Pygmies | -0.012839 | 0.000153 | -84.031 |
| Kuwait-B | Kuwait-P | Anatolia_MLBA | Mbuti_Pygmies | -0.012436 | 0.000148 | -84.035 |
| Kuwait-B | Kuwait-P | Anatolia_N | Mbuti_Pygmies | -0.012278 | 0.000145 | -84.744 |
| Kuwait-B | Kuwait-P | Iran_ChL | Mbuti_Pygmies | -0.01262 | 0.000146 | -86.155 |
| Kuwait-B | Kuwait-P | Europe_EN | Mbuti_Pygmies | -0.012355 | 0.000143 | -86.614 |
| Kuwait-B | Kuwait-P | IranTuran_BA | Mbuti_Pygmies | -0.013005 | 0.000145 | -89.681 |
| Kuwait-B | Kuwait-P | SouthAsia_H | Mbuti_Pygmies | -0.012351 | 0.000136 | -90.567 |
| Kuwait-B | Kuwait-P | Europe_LNBA | Mbuti_Pygmies | -0.012855 | 0.000139 | -92.308 |
| Kuwait-B | Kuwait-P | Steppe_MLBA | Mbuti_Pygmies | -0.012957 | 0.00014 | -92.503 |
| Kuwait-B | Kuwait-P | Steppe_EMBA | Mbuti_Pygmies | -0.013235 | 0.000143 | -92.656 |
| Kuwait-B | Kuwait-P | SPGT | Mbuti_Pygmies | -0.012574 | 0.000134 | -93.782 |

| Pop1 | Pop2 | Pop3 | Pop4 | D val | std err | Z val |
|---|---|---|---|---|---|---|
| Kuwait-B | Kuwait-P | BMAC | Mbuti_Pygmies | -0.012894 | 0.000136 | -94.727 |
| Kuwait-B | Kuwait-S | Iran_HotuIIIb | Mbuti_Pygmies | -0.010315 | 0.00031 | -33.271 |
| Kuwait-B | Kuwait-S | Ust_Ishim | Mbuti_Pygmies | -0.008655 | 0.00017 | -50.929 |
| Kuwait-B | Kuwait-S | Iberia_BA | Mbuti_Pygmies | -0.01201 | 0.000225 | -53.48 |
| Kuwait-B | Kuwait-S | MA1 | Mbuti_Pygmies | -0.010145 | 0.000188 | -54.049 |
| Kuwait-B | Kuwait-S | Kostenki14 | Mbuti_Pygmies | -0.010133 | 0.000175 | -57.824 |
| Kuwait-B | Kuwait-S | Clovis_Anzick | Mbuti_Pygmies | -0.009666 | 0.000166 | -58.286 |
| Kuwait-B | Kuwait-S | Anatolia_ChL | Mbuti_Pygmies | -0.011936 | 0.000198 | -60.167 |
| Kuwait-B | Kuwait-S | Anatolia_Ottoman | Mbuti_Pygmies | -0.010556 | 0.000175 | -60.244 |
| Kuwait-B | Kuwait-S | Iran_N | Mbuti_Pygmies | -0.01013 | 0.000167 | -60.729 |
| Kuwait-B | Kuwait-S | Natufian | Mbuti_Pygmies | -0.011771 | 0.000194 | -60.782 |
| Kuwait-B | Kuwait-S | WestSiberia_HG | Mbuti_Pygmies | -0.010348 | 0.000165 | -62.533 |
| Kuwait-B | Kuwait-S | Switzerland_HG | Mbuti_Pygmies | -0.011098 | 0.000174 | -63.669 |
| Kuwait-B | Kuwait-S | Yamnaya_EBA | Mbuti_Pygmies | -0.011056 | 0.000173 | -63.784 |
| Kuwait-B | Kuwait-S | Iran_recent | Mbuti_Pygmies | -0.011542 | 0.000181 | -63.865 |
| Kuwait-B | Kuwait-S | Steppe_IA | Mbuti_Pygmies | -0.011064 | 0.000173 | -63.952 |
| Kuwait-B | Kuwait-S | Indus_Diaspora | Mbuti_Pygmies | -0.009857 | 0.000154 | -63.976 |
| Kuwait-B | Kuwait-S | Anatolia_IA | Mbuti_Pygmies | -0.01143 | 0.000177 | -64.575 |
| Kuwait-B | Kuwait-S | EHG | Mbuti_Pygmies | -0.010785 | 0.000163 | -66.18 |
| Kuwait-B | Kuwait-S | Turkmenistan_IA | Mbuti_Pygmies | -0.011062 | 0.000166 | -66.652 |
| Kuwait-B | Kuwait-S | IranTuran_N | Mbuti_Pygmies | -0.010278 | 0.000149 | -68.811 |
| Kuwait-B | Kuwait-S | CHG | Mbuti_Pygmies | -0.010743 | 0.000155 | -69.093 |
| Kuwait-B | Kuwait-S | Namazga_CA | Mbuti_Pygmies | -0.010547 | 0.000152 | -69.43 |
| Kuwait-B | Kuwait-S | SHG | Mbuti_Pygmies | -0.011086 | 0.000155 | -71.497 |
| Kuwait-B | Kuwait-S | Levant_N | Mbuti_Pygmies | -0.012051 | 0.000166 | -72.791 |
| Kuwait-B | Kuwait-S | WHG | Mbuti_Pygmies | -0.011186 | 0.000152 | -73.634 |
| Kuwait-B | Kuwait-S | Steppe_LBA | Mbuti_Pygmies | -0.010802 | 0.000146 | -74.236 |
| Kuwait-B | Kuwait-S | Armenia_ChL | Mbuti_Pygmies | -0.011618 | 0.000153 | -76.005 |
| Kuwait-B | Kuwait-S | Armenia_EBA | Mbuti_Pygmies | -0.011519 | 0.00015 | -76.776 |
| Kuwait-B | Kuwait-S | Levant_BA | Mbuti_Pygmies | -0.012209 | 0.000159 | -76.866 |
| Kuwait-B | Kuwait-S | IranTuran_BA | Mbuti_Pygmies | -0.010751 | 0.00014 | -77.054 |
| Kuwait-B | Kuwait-S | SouthAsia_H | Mbuti_Pygmies | -0.010246 | 0.00013 | -78.556 |
| Kuwait-B | Kuwait-S | Anatolia_EBA | Mbuti_Pygmies | -0.011735 | 0.000149 | -78.749 |
| Kuwait-B | Kuwait-S | Iran_ChL | Mbuti_Pygmies | -0.011279 | 0.000143 | -79.135 |
| Kuwait-B | Kuwait-S | Anatolia_MLBA | Mbuti_Pygmies | -0.011904 | 0.000147 | -80.929 |
| Kuwait-B | Kuwait-S | SPGT | Mbuti_Pygmies | -0.010397 | 0.000128 | -81.142 |
| Kuwait-B | Kuwait-S | Steppe_EMBA | Mbuti_Pygmies | -0.011055 | 0.000133 | -82.933 |
| Kuwait-B | Kuwait-S | Steppe_MLBA | Mbuti_Pygmies | -0.011265 | 0.000133 | -84.545 |
| Kuwait-B | Kuwait-S | BMAC | Mbuti_Pygmies | -0.010909 | 0.000128 | -85.343 |
| Kuwait-B | Kuwait-S | Europe_LNBA | Mbuti_Pygmies | -0.011533 | 0.000134 | -85.785 |
| Kuwait-B | Kuwait-S | Anatolia_N | Mbuti_Pygmies | -0.012307 | 0.000141 | -87.182 |
| Kuwait-B | Kuwait-S | Europe_EN | Mbuti_Pygmies | -0.012301 | 0.00014 | -87.6 |
| Kuwait-P | Kuwait-S | SPGT | Mbuti_Pygmies | 0.002178 | 0.000069 | 31.616 |

| Pop1 | Pop2 | Pop3 | Pop4 | D val | std err | Z val |
|---|---|---|---|---|---|---|
| Kuwait-P | Kuwait-S | SouthAsia_H | Mbuti_Pygmies | 0.002105 | 0.00007 | 29.992 |
| Kuwait-P | Kuwait-S | Steppe_EMBA | Mbuti_Pygmies | 0.00218 | 0.000075 | 29.145 |
| Kuwait-P | Kuwait-S | IranTuran_BA | Mbuti_Pygmies | 0.002254 | 0.00008 | 28.318 |
| Kuwait-P | Kuwait-S | BMAC | Mbuti_Pygmies | 0.001985 | 0.00007 | 28.159 |
| Kuwait-P | Kuwait-S | IranTuran_N | Mbuti_Pygmies | 0.00256 | 0.000095 | 26.937 |
| Kuwait-P | Kuwait-S | Indus_Diaspora | Mbuti_Pygmies | 0.002369 | 0.000089 | 26.628 |
| Kuwait-P | Kuwait-S | WestSiberia_HG | Mbuti_Pygmies | 0.002704 | 0.000103 | 26.236 |
| Kuwait-P | Kuwait-S | Namazga_CA | Mbuti_Pygmies | 0.002245 | 0.00009 | 25.018 |
| Kuwait-P | Kuwait-S | Steppe_MLBA | Mbuti_Pygmies | 0.001692 | 0.000072 | 23.454 |
| Kuwait-P | Kuwait-S | CHG | Mbuti_Pygmies | 0.002226 | 0.000096 | 23.284 |
| Kuwait-P | Kuwait-S | Steppe_LBA | Mbuti_Pygmies | 0.001912 | 0.000083 | 23.109 |
| Kuwait-P | Kuwait-S | Iran_N | Mbuti_Pygmies | 0.002393 | 0.000105 | 22.777 |
| Kuwait-P | Kuwait-S | EHG | Mbuti_Pygmies | 0.002293 | 0.000103 | 22.195 |
| Kuwait-P | Kuwait-S | MA1 | Mbuti_Pygmies | 0.00254 | 0.000118 | 21.499 |
| Kuwait-P | Kuwait-S | Clovis_Anzick | Mbuti_Pygmies | 0.00208 | 0.000104 | 19.986 |
| Kuwait-P | Kuwait-S | Yamnaya_EBA | Mbuti_Pygmies | 0.00212 | 0.000112 | 19.008 |
| Kuwait-P | Kuwait-S | Europe_LNBA | Mbuti_Pygmies | 0.001322 | 0.000073 | 18.046 |
| Kuwait-P | Kuwait-S | SHG | Mbuti_Pygmies | 0.001616 | 0.000091 | 17.773 |
| Kuwait-P | Kuwait-S | Steppe_IA | Mbuti_Pygmies | 0.001838 | 0.000106 | 17.397 |
| Kuwait-P | Kuwait-S | Turkmenistan_IA | Mbuti_Pygmies | 0.001812 | 0.000112 | 16.204 |
| Kuwait-P | Kuwait-S | Iran_ChL | Mbuti_Pygmies | 0.001341 | 0.000086 | 15.528 |
| Kuwait-P | Kuwait-S | Armenia_EBA | Mbuti_Pygmies | 0.00133 | 0.000089 | 14.906 |
| Kuwait-P | Kuwait-S | Anatolia_Ottoman | Mbuti_Pygmies | 0.001442 | 0.000105 | 13.797 |
| Kuwait-P | Kuwait-S | Armenia_ChL | Mbuti_Pygmies | 0.001115 | 0.000086 | 12.907 |
| Kuwait-P | Kuwait-S | WHG | Mbuti_Pygmies | 0.001122 | 0.000097 | 11.572 |
| Kuwait-P | Kuwait-S | Ust_Ishim | Mbuti_Pygmies | 0.001127 | 0.000106 | 10.661 |
| Kuwait-P | Kuwait-S | Switzerland_HG | Mbuti_Pygmies | 0.001212 | 0.000114 | 10.658 |
| Kuwait-P | Kuwait-S | Iran_HotuIIIb | Mbuti_Pygmies | 0.002132 | 0.000205 | 10.382 |
| Kuwait-P | Kuwait-S | Iran_recent | Mbuti_Pygmies | 0.001099 | 0.000115 | 9.591 |
| Kuwait-P | Kuwait-S | Kostenki14 | Mbuti_Pygmies | 0.001004 | 0.00011 | 9.164 |
| Kuwait-P | Kuwait-S | Anatolia_IA | Mbuti_Pygmies | 0.000904 | 0.000109 | 8.283 |
| Kuwait-P | Kuwait-S | Anatolia_MLBA | Mbuti_Pygmies | 0.000532 | 0.000083 | 6.436 |
| Kuwait-P | Kuwait-S | Anatolia_ChL | Mbuti_Pygmies | 0.000799 | 0.000125 | 6.382 |
| Kuwait-P | Kuwait-S | Anatolia_EBA | Mbuti_Pygmies | 0.000484 | 0.000088 | 5.507 |
| Kuwait-P | Kuwait-S | Iberia_BA | Mbuti_Pygmies | 0.000622 | 0.000142 | 4.391 |
| Kuwait-P | Kuwait-S | Europe_EN | Mbuti_Pygmies | 0.000054 | 0.000077 | 0.694 |
| Kuwait-P | Kuwait-S | Anatolia_N | Mbuti_Pygmies | -0.000029 | 0.000077 | -0.374 |
| **Kuwait-P** | **Kuwait-S** | **Levant_BA** | **Mbuti_Pygmies** | **-0.000746** | **0.000096** | **-7.792** |
| **Kuwait-P** | **Kuwait-S** | **Levant_N** | **Mbuti_Pygmies** | **-0.001042** | **0.000098** | **-10.645** |
| **Kuwait-P** | **Kuwait-S** | **Natufian** | **Mbuti_Pygmies** | **-0.002034** | **0.000115** | **-17.622** |
